# Genetic Disruption of Serine Biosynthesis is a Key Driver of Macular Telangiectasia Type 2 Etiology and Progression

**DOI:** 10.1101/2020.02.04.934356

**Authors:** Roberto Bonelli, Brendan R E Ansell, Luca Lotta, Thomas Scerri, Traci E Clemons, Irene Leung, The MacTel Consortium, Tunde Peto, Alan C Bird, Ferenc B Sallo, Claudia Langenberg, Melanie Bahlo

**Affiliations:** Department of Medical Biology, The University of Melbourne, 3052, Parkville, Victoria, Australia; Population Health and Immunity Division, Walter and Eliza Hall Institute of Medical Research, 3052, Parkville, Victoria, Australia; MRC Epidemiology Unit, University of Cambridge, CB2 0SL, Cambridge, UK.; The EMMES Corporation, Rockville, 20850, Maryland, United States; Department of Research and Development, Moorfields Eye Hospital NHS Foundation Trust, EC1V 2PD, London, United Kingdom; A list of members and affiliations is provided in Table S8; Department of Ophthalmology, Queen’s University, Belfast, BT7 1NN, United Kingdom; Inherited Eye Disease, Moorfields Eye Hospital NHS Foundation Trust, EC1V 2PD, London, United Kingdom; Department of Ophthalmology, University of Lausanne, Hôpital Ophtalmique Jules-Gonin, Fondation Asile des aveugles, Switzerland

**Keywords:** Retinal Disease, Mendelian Randomization, Metabolomics, GWAS, Serine

## Abstract

**Purpose:** Macular telangiectasia type 2 (MacTel) is a rare, heritable and largely untreatable retinal disorder, often comorbid with diabetes. Genetic risk loci subtend retinal vascular calibre, and glycine/serine/threonine metabolism genes. Serine deficiency may contribute to MacTel via neurotoxic deoxysphingolipid production, however, an independent vascular contribution is also suspected. Here we use statistical genetics to dissect the causal mechanisms underpinning this complex disease.

**Methods:** We integrated genetic markers for MacTel, vascular, and metabolic traits, and applied Mendelian randomization, MTAG, and conditional/interaction genome-wide association analysis to discover causal contributors to both disease, and spatial retinal imaging sub-phenotypes.

**Results:** Serine was a key causal driver of disease occurrence and progression, with a lesser contribution to type 2 diabetes risk. Conversely, glycine, threonine and retinal vascular traits are unlikely to be causal for MacTel. Conditional regression analysis resolved three novel disease loci independent of endogenous serine biosynthetic capacity. By aggregating retinal phenotypes into endophenotypes, we demonstrate that SNPs constituting independent risk loci act via related endophenotypes.

**Discussion:** Our findings will aid in early diagnosis and accurate prognosis of MacTel, and improve prospects for effective therapeutic intervention. Our integrative genetics approach also serves as a useful template for post-GWAS analyses in other disorders.

## Introduction

### MacTel Disease and previous GWAS study

Macular telangiectasia type II (MacTel^1^), is a rare degenerative eye disease affecting the macula^2, 3^. MacTel is bilateral and progressively affects visual acuity^4, 5^, reading ability^6^, and vision-related quality of life^7, 8^. A successful phase II clinical trial has recently been reported for an intravitreal encapsulated cell therapy implant, showing efficacy in slowing disease progression^9^. However, less invasive, non-surgical and more economical therapies are lacking. Given the rarity of MacTel and its subtle clinical signs, requiring several ophthalmological diagnostic methods, MacTel has been largely under/misdiagnosed^4^. Hence further insight into the genetic basis of MacTel is key for accurate diagnosis, identifying future therapies, developing predictive models for the disease, and increasing prognostic accuracy.

We previously published the first genome-wide association study on 476 MacTel patients and 1733 controls^10^, identifying and replicating five loci. A single nucleotide polymorphism (SNP) at locus 5q14.3 (rs73171800) showed the strongest association with the disease and was previously identified to be associated with the quantitative traits of retinal venular and arterial calibre^11, 12^. The other four loci, 1p12 (rs477992), 2q34 (rs715), 7p11.2 (rs4948102), and 3q21.3 (rs9820286) were implicated in glycine and serine metabolism^13, 14^. Importantly, loci 3q21.3 and 7p11.2 did not reach genome-wide significance, and the SNPs at locus 3q21.3 were only in proximity, but not in linkage disequilibrium (LD) with those associated with MacTel. We additionally measured and found significant depletion of serum serine, glycine and threonine in MacTel patients compared to controls. These data provided the first insight into the genetic complexity underpinning MacTel, and highlighted the potential involvement of metabolic and vascular trait disturbances in disease aetiology.

Glycine and serine are involved in many fundamental biochemical reactions. Pathogenic variants in genes involved in serine and glycine synthesis lead to severe congenital disorders such as phosphoglycerate dehydrogenase deficiency (*PHGDH*, <u>601815</u>) and glycine encephalopathy (*GLDC*, <u>605899</u>). Glycine and serine can be synthesized from one another, from other metabolic compounds or obtained through dietary intake.

A recent study reported the co-occurrence of MacTel in patients with a rare neuropathy, hereditary sensory and autonomic systemic neuropathy type I, or HSAN1 (<u>162400</u>)^15^. This neuropathy results from pathogenic loss of function variants in the enzyme serine palmityltransferase, leading to accumulation of neurotoxic deoxysphingolipids. A similar accumulation of deoxysphingolipids was demonstrated in MacTel patients, but unlike HSAN1, was attributed to systemic serine deficiency.

MacTel is often comorbid with type 2 diabetes (T2D)^16^. It is unknown whether traits such as metabolite abundances, T2D, or retinal vascular calibre, now known to be associated with MacTel, are causative or merely associated with the disease for other reasons. Causation hypotheses are now routinely investigated using Mendelian Randomization (MR). MR borrows from Instrumental Variable (IV) analysis and assumes that if a disease is caused by a particular intermediate phenotype, and the latter is caused by genetic variants, then the same genetic variants should be also associated with the disease^17–19^. Genetic variants involved in retinal vascular calibre traits and T2D have been previously identified^11, 12, 20^ and the largest metabolomic GWAS meta-analysis to date recently identified almost 500 loci that are associated with differences in 142 serum metabolite levels in humans^21, 22^. The results from these studies can be used to investigate causal drivers of MacTel susceptibility via the MR approach.

GWA studies on rare complex traits such as MacTel are inevitably underpowered due to the difficulty in recruiting a large number of samples. Methods leveraging on well-powered and publicly available GWAS have now been developed. An example is Multi-Trait Association GWAS (MTAG)^23^ which exploits the genetic correlation between traits to identify new loci.

Progress in MacTel diagnosis and treatment is complicated by a lack of clear indicators of early stages and by the substantial inter-patient and interocular heterogeneity of retinal phenotypes. Apportioning genetic variation to separate retinal malformations has the potential to shed light on the different biological mechanisms via which each locus contributes to the disease. Performing joint genetic and retinal phenotypic data thus investigates whether different retinal abnormalities are consequences of common or largely independent biological perturbations.

In this study, we test for the causality of several traits on MacTel disease (**Figure 1A**). We exploit independent studies on MacTel-related traits to increase discovery power in our disease cohort, and both identify new disease loci while further resolving previously identified loci (**Figure 1B**). We use extensive retinal phenotypic data from MacTel patients to identify key genetic drivers of specific subphenotypes (**Figure 1C**). We collate genetic loci into functional groups by testing their combined effects on ocular spatial phenotypes (**Figure 1D**). This study represents the first integrated analysis of MacTel genotypic data with MacTel-related traits as well as retinal imaging phenotypic data. Our study serves as a model for post-GWAS studies with phenotyping datasets.

**Figure 1.**
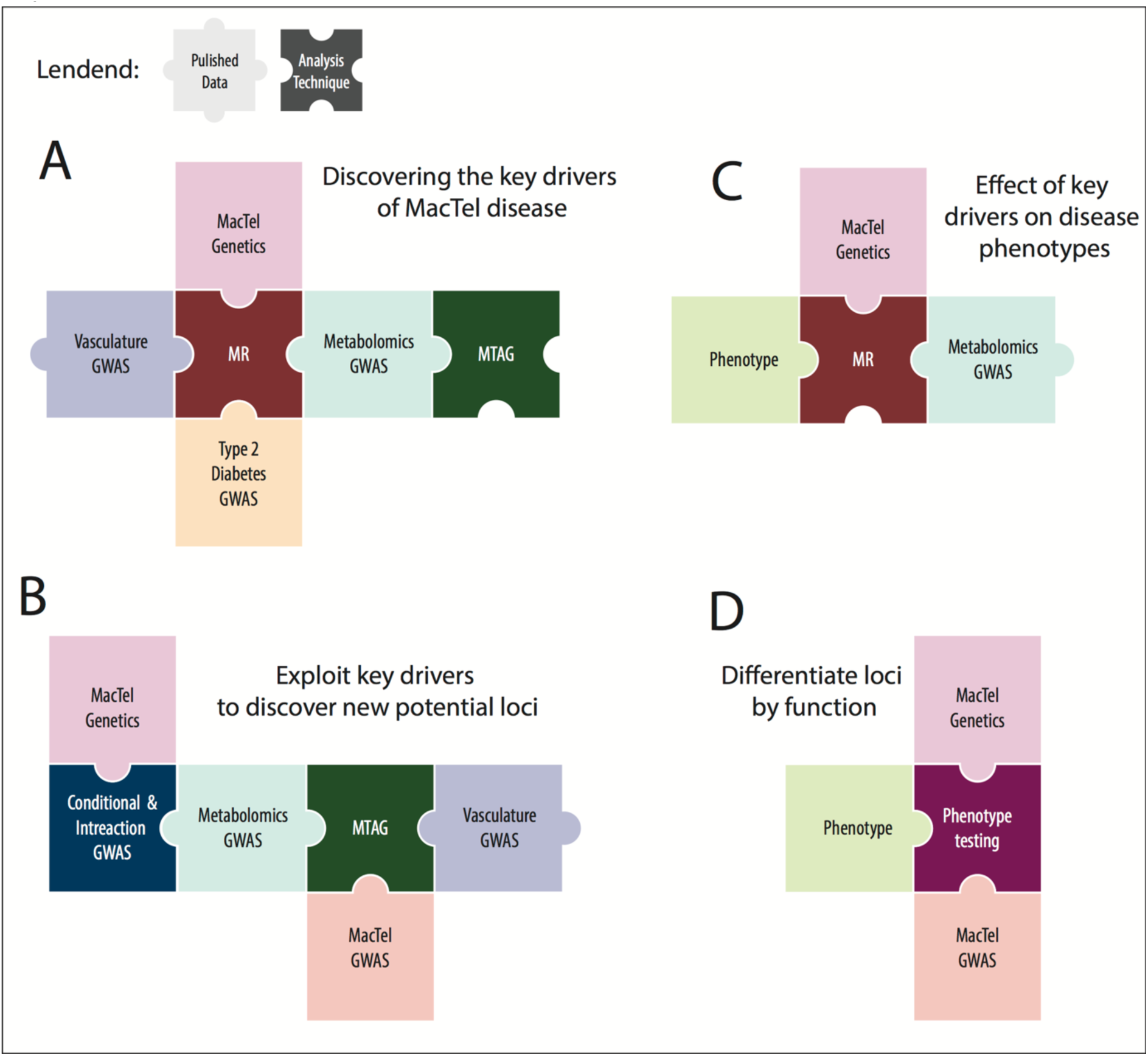
Study conceptual map. Each panel represents a study aim and depicts the data and the analysis technique used. Data is presented as pastel color pieces with black typing while analysis techniques are presented as darker panels with white writing. MacTel genetics refers to individual level SNP data available from our previous GWAS study. MacTel GWAS refers to the MacTel GWAS summary statistics. Vasculature GWAS refers to the summary statistics data available from two previous retinal vasculature calibre genetic studies^11, 12^. Metabolomics GWAS refers to summary statistics data available from a recently published metabolomics GWAS^21, 22^. T2D GWAS refers to summary statistics data available from GWAS study on T2D^20^. MTAG: Multi-Trait Association GWAS; MR: Mendelian Randomization.

## Material and Methods

Unless otherwise stated, all statistical and computational analyses were performed using the R statistical software, version 3.5.1. Corrected p-values less than 0.05 were considered statistically significant. For genome-wide association analyses, uncorrected p-values less than 5e-8 were considered genome-wide significant.

### DNA samples and SNP genotyping

Genotypic data was available for 476 MacTel patients and 1733 controls. SNPs were genotyped using the Illumina 5.0M chip as described previously^10^. Retinal phenotypic data was available for 455 patients MacTel patients. Genetic predictors for metabolites and retinal vascular calibre, were provided by the corresponding authors of each study^11, 12, 21, 22^.

### Mendelian Randomization Procedures

To perform Mendelian randomization analysis on MacTel with metabolites, T2D and retinal vasculature we used the allele-score method for individual-level genetic data as described by Burgess et al. 2016^24^. Using a p-value threshold of 5e-8 we extracted SNPs significantly associated with serum metabolite concentrations or other traits of interest (retinal venular calibre, retinal arteriolar calibre, and T2D). We estimated Genetically Predicted Metabolites (GPMs) and Genetically Predicted Traits (GPTs) (**Table S1**) for each subject by using the SNPs magnitudes as weights, in the manner of a polygenic risk score (**Supplementary Methods**). Using logistic regression models we tested for association between GPMs, or GPTs and MacTel susceptibility. This analysis was corrected for genetically determined sex at birth and the first principal component to account for batch effects including population stratification, as in our previous publication^10^. We used Benjamini-Hochberg multiple testing correction to control the false discovery rate threshold of 5%. Conditional modelling approach was used to identify GPMs independently associated with the disease. Specifically, the most significant GPM was iteratively added as covariate in the regression model until no GPM was significant after FDR correction.

**Table 1.** Significant associations between GPMs and GPT and MacTel. Regression coefficients are presented in the “Beta” column. Odds Ratios (OR) are relative to a single standard deviation increase on the GPM/GPT scale. Odds Ratios 95% confidence intervals are shown in the column 95% C.. Benjamini-Hochberg adjusted P-values are presented under the column P-value.

| GP Metabolite / Trait | Beta | OR | 95% C.I. | P-value | Biochemical Family |
| --- | --- | --- | --- | --- | --- |
| Serine | -0.65 | 0.52 | 0.46 - 0.58 | 8.38E-28 | Amino Acids and Biogenic Amines |
| Glycine | -0.58 | 0.56 | 0.50 - 0.63 | 5.44E-19 | Amino Acids and Biogenic Amines |
| lysoPhosphatidylcholine acyl<br>C14:0 | 0.21 | 1.23 | 1.11 - 1.37 | 2.48E-03 | Glycerophospholipids |
| Phosphatidylcholine diacyl<br>C32:1 | 0.21 | 1.23 | 1.11 - 1.37 | 2.48E-03 | Glycerophospholipids |
| Phosphatidylcholine diacyl<br>C34:1 | 0.21 | 1.23 | 1.11 - 1.37 | 2.48E-03 | Glycerophospholipids |
| Phosphatidylcholine with acyl-<br>alkyl residue C38:1 | 0.20 | 1.22 | 1.10 - 1.36 | 2.97E-03 | Glycerophospholipids |
| Arginine | 0.17 | 1.18 | 1.07 - 1.31 | 2.83E-02 | Amino Acids and Biogenic Amines |
| Phenylalanine | -0.17 | 0.85 | 0.76 - 0.94 | 2.85E-02 | Amino Acids and Biogenic Amines |
| Arteriolar Retinal Calibre | 0.32 | 1.38 | 1.25 - 1.52 | 2.22E-10 | Vasculature Trait |
| Venular Retinal Calibre | 0.18 | 1.19 | 1.08 - 1.32 | 7.00E-04 | Vasculature Trait |
| T2D | 0.17 | 1.19 | 1.07 - 1.33 | 7.72E-04 | Comorbidity |

### Conditional and Interaction GWAS and MTAG analysis

Conditional GWAS were performed by including GPMs or GPTs in the logistic regression models as covariates. For the interaction GWAS, an interaction term was included for each SNP and the GPM or GPT combination of interest. Each model also included genetically determined sex at birth and the first principal component^10^. Analyses were performed using Plink v1.9^25^. Results from the conditional and interaction GWAS were analysed using FUMA^26^. LocusZoom plots were produced using the LocusZoom software ^27^. MTAG analysis was performed by integrating the available summary statistics for each trait of interest using the MTAG software^23^.

### Retinal phenotypes

We used longitudinal retinal phenotypic data on 1,716 patients (3,410 eyes) collected from Natural History Observation and Registry studies of Macular Telangiectasia^28^. The data consisted of 143 spatial measurements of retinal phenotypes (**Table S2)**. A detailed description of the corresponding methods can be found elsewhere^28^. Each phenotype was measured in 9 different sub-fields of the retina (defined by the ETDRS grid^29^ **Figure S1**). The phenotype data was cleaned by performing missing data imputation (**Supplementary Methods**). The cleaned dataset contained 119 phenotypes that were collapsed into 30 biologically relevant endophenotypes using factor analysis (**Supplementary Methods**).

**Table 2.** Genome-wide significant loci for MacTel from MTAG analysis using retinal venular (Ven) and arteriolar (Art) calibre, serine (Ser) and glycine (Gly). Novel genome-wide significant loci are identified by the column New GW. Bold text indicates genome-wide significant P-values on respective original studies. Grey italicised p-values are associations expected to be null *a priori*.

| rsid | Locus | Candidate<br>Gene | MTAG P-val<br>MacTel | New<br>GW. | P-val Art<br>Sim et al<br>2013 | P-val Ven<br>Ikram et al<br>2010 | P-val Gly<br>Lotta et al<br>2019 | P-val Ser<br>Lotta et<br>al 2019 | P-val<br>MacTel<br>Scerri et<br>al 2017 |
| --- | --- | --- | --- | --- | --- | --- | --- | --- | --- |
| rs3815630 | 2q34 | *CPS1* | **7.62E-143** | No | *1.51E-01* | *6.23E-01* | **2.23E-308** | **6.50E-31** | **4.43E-08** |
| rs1035387 | 5q14.3 | *TMEM161B-*<br>*AS1* | **1.16E-16** | No | **1.50E-09** | **1.15E-08** | *5.43E-01* | *7.53E-01* | **1.16E-16** |
| rs477992 | 1p12 | *PHGDH* | **2.21E-15** | No | *7.44E-01* | *7.95E-04* | **6.09E-07** | **2.01E-80** | **2.60E-12** |
| rs4543497 | 7p11.2 | *PSPH* | **2.22E-15** | Yes | *3.56E-01* | *2.07E-01* | **9.01E-28** | **3.29E-54** | 8.11E-06 |
| rs4841132 | 8p23.1 | *LOC157273* | **8.43E-14** | Yes | *5.55E-02* | *2.95E-02* | **1.73E-34** | 1.23E-04 | 4.54E-02 |
| rs1563075 | 16q23 | *GCSH* | **1.79E-13** | Yes | *6.86E-01* | *3.77E-01* | **2.57E-28** | 1.19E-01 | 4.66E-03 |
| rs2954021 | 8q24.1 | *TRIB1* | **1.53E-12** | Yes | *3.89E-01* | *6.33E-01* | **2.35E-21** | **2.40E-10** | 8.70E-05 |
| rs4742212 | 9p24 | *GLDC* | **1.19E-09** | Yes | *4.73E-01* | *6.88E-01* | **3.06E-34** | 1.69E-04 | 2.82E-01 |

### Investigating the relationship of retinal endophenotypes with genetic drivers

We tested for association between retinal endophenotypes and significant disease-associated GPMs, GPTs and previously prioritised SNPs. The dataset containing retinal endophenotypes and genetic information included 3,280 observations from 455 MacTel patients (907 eyes in total) with an average of 3.6 observations per eye over 10 years. Association testing was performed using a linear mixed model approach, assuming an additive effect of SNP alleles that increased MacTel risk. P-values were corrected using an adaptive Benjamini-Hochberg procedure^30^. Further details are provided in **Supplementary Methods**.

## Results

### Discovering key drivers of MacTel disease: Metabolites

The MR procedure (**Figure 1A**) applied to the metabolite panel highlighted 14 GP metabolites significantly associated with MacTel (**Table 1, Table S3)**. The two most significant metabolites were GP serine and GP glycine In contrast to the metabolomics results from our previous metabolomics study with directly measured metabolites, GP threonine was not significant. Other significant metabolites included GP arginine, GP phenylalanine and GP phosphatidylcholine species. GP alanine was borderline significant.

The significant GPMs correlated with each other in our sample (**Figure S2**), mirroring correlations observed with directly measured metabolite abundances. When all GPMs were re-tested in a model including GP serine as a covariate, GP glycine was the only associated metabolite (**Table S3**) suggesting that the effect of GP glycine on MacTel is largely due to biochemical interactions with serine. GP serine remained significant when included alongside GP glycine. No other GP metabolites remained significant once GP serine and GP glycine had been included (**Table S3**).

The GWAS used to predict the genetic complement of serine was underpowered compared to the glycine GWAS^22^ (**Supplementary Materials**). However, biochemically-based genetic correlation exists between the two. We attempted to boost the power of the serine GWAS results by combining them with the glycine GWAS results in an MTAG analysis (**Figure 1A**). The newly identified MTAG serine loci were used to construct a new and more powerful genetic predictor of serine (GP MTAG serine, **Table S1**). We found that GP MTAG serine encompassed the entire metabolic predictive signal for MacTel, as the original GP glycine was no longer required in the model once GP MTAG serine was included. Further MR testing of a broader set of 248 GP metabolite abundances using results from Shin et al^14, 31^ (**Table S1**) found a few additional significant univariate contribution from other metabolites (**Supplementary Materials** and **Table S4**). However, none remained significant after inclusion of GP serine as a covariate.

### Discovering key drivers of MacTel disease: T2D and retinal vasculature

Genetically predicted T2D was significantly associated with MacTel, as were both GP arteriolar and venular calibre when tested separately. However, the vasculature trait results were driven by a single SNPs: rs2194025 (for arteriolar calibre) and rs17421627 (for venular calibre), both in strong LD with SNP rs73171800 at 5q14.3 previously identified in our original GWAS^10^. When including both vascular traits in the same model, only arteriolar calibre remained significant, due to the reduced number of SNPs (N = 2) used to construct that GPT (**Figure S3**). When GPTs for retinal arteriolar calibre and T2D were included in a model with GP glycine and serine, all remained significant.

As a quality control analysis, we compared the effect sizes for each instrumental SNP on both MacTel and the metabolites or traits to inspect for potential SNP outlier effects of pleiotropy (**Figure 2**). A negative, linear relationship was detected between the effect on metabolic abundance and MacTel risk for the SNPs used to define glycine and serine. A positive relationship was observed for T2D and vasculature traits, while no relationship was detected for threonine which was used as a control trait, where no significant association was observed. We additionally observed that the regression line for all significant traits except the vasculature related traits, did not have intercept terms that were significantly different from 0 (b_0,*serine*_= – 0.07, p=0.2; b_0,*glycine*_=0.070, p=0.131; b_0,*T2D*_=0.002, p=0.88) suggesting a non-pleiotropic effect of the instruments used to test such traits^32^. The same conclusion was drawn when Glycine and T2D SNPs considered to have potential outlier effects were removed.

**Figure 2.**
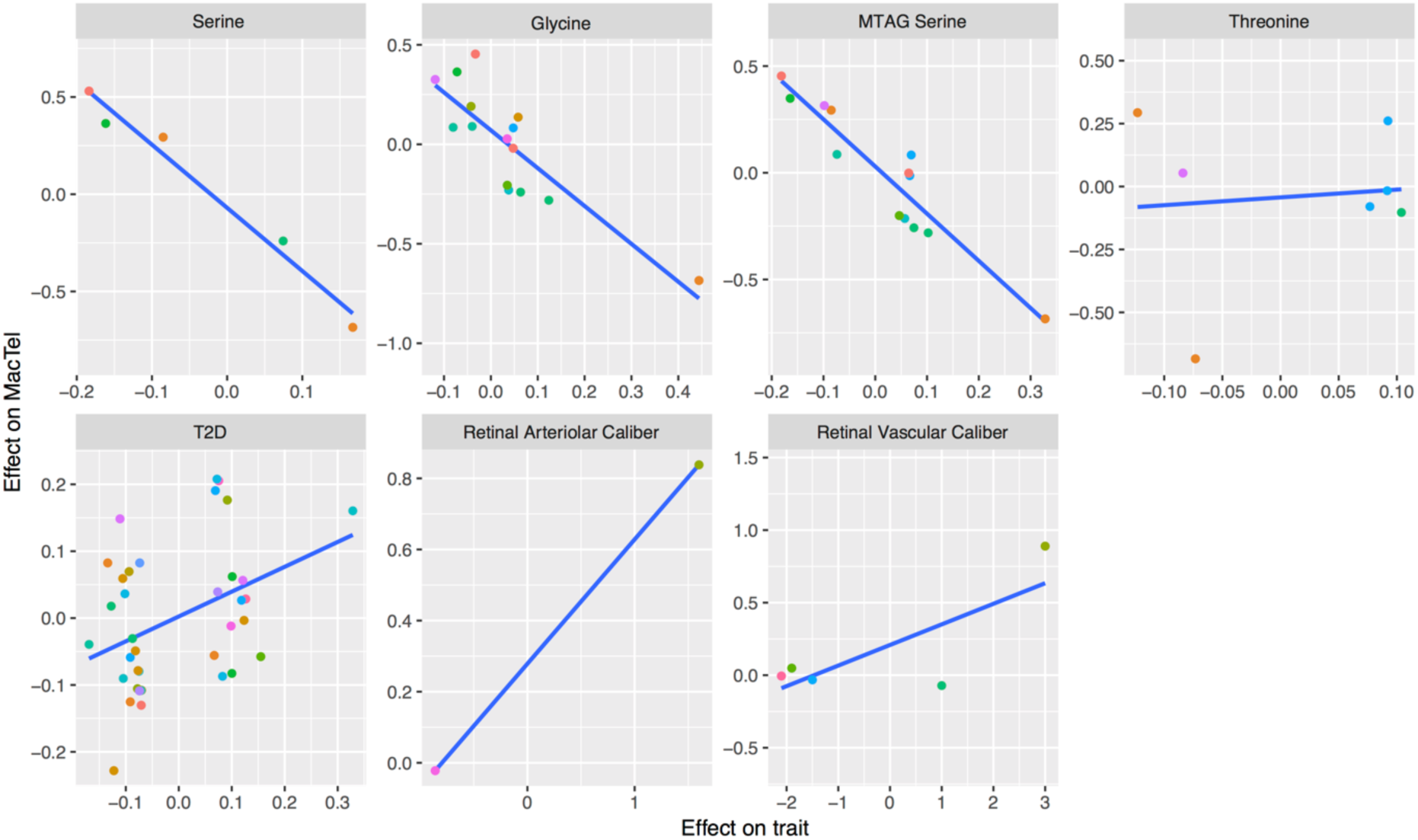
Effect size comparisons between MacTel and specific trait of SNPs used to define GPM and GPT. Each dot represents a SNP. Each panel represents a trait and contains the SNPs that were found to have a genome-wide significant effect on that trait. The x-axis captures the effect sizes of the SNPs on the trait while the y-axis captures the effect of the same SNPs on MacTel risk. The blue line represents a simple regression line between the two. Metabolites or traits causally affecting the disease are expected to have correlated effect sizes. Colour of the dots represents chromosomal positions where different chromosomes are represented by different colours.

### Using causal traits to discover new disease susceptibility loci for MacTel

#### MTAG results

To discover new loci putatively involved in MacTel, we performed MTAG analysis by including GWAS summary statistics for glycine, serine, retinal venular calibre, and retinal arteriolar calibre along-side MacTel (**Figure 1B**). T2D was not included as genome-wide summary statistics were not available. The number of SNPs shared across all datasets was 799,436 (maximal FDR = 0.397). This analysis confirmed the three previously identified MacTel risk loci^10^ and revealed 5 novel loci attaining genome-wide significance (**Table 2, Table S5** for details).

Among the five new genome-wide significant loci was locus 7p11.2 which did not reach genome-wide significance in the original GWAS but was deemed important given its established relationship with glycine and serine^10^. We did not observe any significant MTAG results for locus 3q21.3, originally proposed to affect MacTel though a connection with glycine and serine^10^. This SNP was in fact not genome-wide significant for either glycine (p = 0.041) or serine (p = 0.65) and is likely to present either a false positive or a GWAS signal not contributed to by glycine or serine.

#### Conditional and Interaction GWAS

To uncover genetic correlates of MacTel independent of glycine and serine, we performed a GWAS conditioning on these genetically predicted traits (**Figure 1B**). This analysis identified four genome-wide significant peaks (**Figure S4 a)**.

As expected, the original signal on locus 5q14.3 (rs73171800) related to vascular calibre, remained significant, confirming the independence of this locus from genetic drivers of serine and glycine. A second genome-wide significant signal on locus 3p24.1 (rs35356316, p = 3.10e-08) was identified and was situated in a ‘gene desert’ proximal to the genes *EOMES* and *SLC4A4*. Another single SNP in very close proximity to rs35356316 reached genome-wide significance as in the original MacTel GWAS study^10^ but was believed to be a false positive, given the lack of LD with any other significant SNP (**Figure S4 b**).

The remaining two conditionally significant SNPs tagged independent signals in locus 19p13.2 (**Figure S4 c-d**). SNP rs36259 is an exonic non-synonymous SNP located in the *CERS4* gene which achieved close to genome-wide significance (p=6.270e-08) and was nominally significant in our original study (p = 1.69e-7). In close proximity, we found an independent intergenic SNP rs4804075 (p = 3.72e-07) which lacked evidence of an eQTL effect on any neighbouring genes, and which did not reach genome-wide significance. These results remain significant when conditioning on SNP rs73171800 (locus 5q14.3) and GP T2D.

We performed additional GWAS analyses testing for interactions between all SNPs with GP serine, glycine, T2D, and SNP rs73171800 (**Figure 1B**). However, no further significant interacting loci were found (**Figure S5 a-d**).

### Effect of key drivers on retinal endophenotypes

Given the small sample size of the endophenotypic data (N = 455), we only tested for association between endophenotypes and GP glycine, serine and T2D, as these bore the clearest association with disease etiology (**Figure 1C, Table S6**). No significant association between GP glycine or T2D and endophenotypes was found after correcting for GP serine, and we, therefore, excluded them from additional testing. High GP serine levels were found to be protective against loss of retinal transparency, leakage at the outer capillary network and the retinal pigment epithelium (RPE) in MacTel progression areas, and leakage at the RPE in the MacTel area. We also found suggestive significance for protection from perivascular pigment clustering in the MacTel area and progression area. Higher GP serine levels were associated with protection from macular thinning in the MacTel area, inferior inner area, and suggestive protection on nasal inner area, and foveal area.

### Determining the effects of GWAS loci on endophenotypes

To determine the likely functional impacts of the original GWAS loci we tested for associations with retinal endophenotypes (**Figure 1D, Figure 3, Table S6**). No associations remained significant after correction for multiple testing due to the modest sample and effect sizes of individual SNPs on disease endophenotypes. Using a nominally (uncorrected) significant p-value threshold denoted p* < 0.05 in an exploratory approach, we observed that locus 1p12 (rs477992) was associated with an increased risk of perivascular pigment clustering in the progression area, but protected against risk of blunted vessels. Furthermore, locus 2q34 (rs715) increased the risk of macular thinning in the MacTel area and nasal inner area and the risk of perivascular pigment clustering in the MacTel area. Finally, the MacTel risk allele at locus 7p11.2 (rs4948102) increased the risk of retinal transparency, leakage at the RPE in both the MacTel area and progression area, as well as presence of inner empty spaces and EZ break. This locus however protected against leakage in the outer layer of the foveal area.

**Figure 3.**
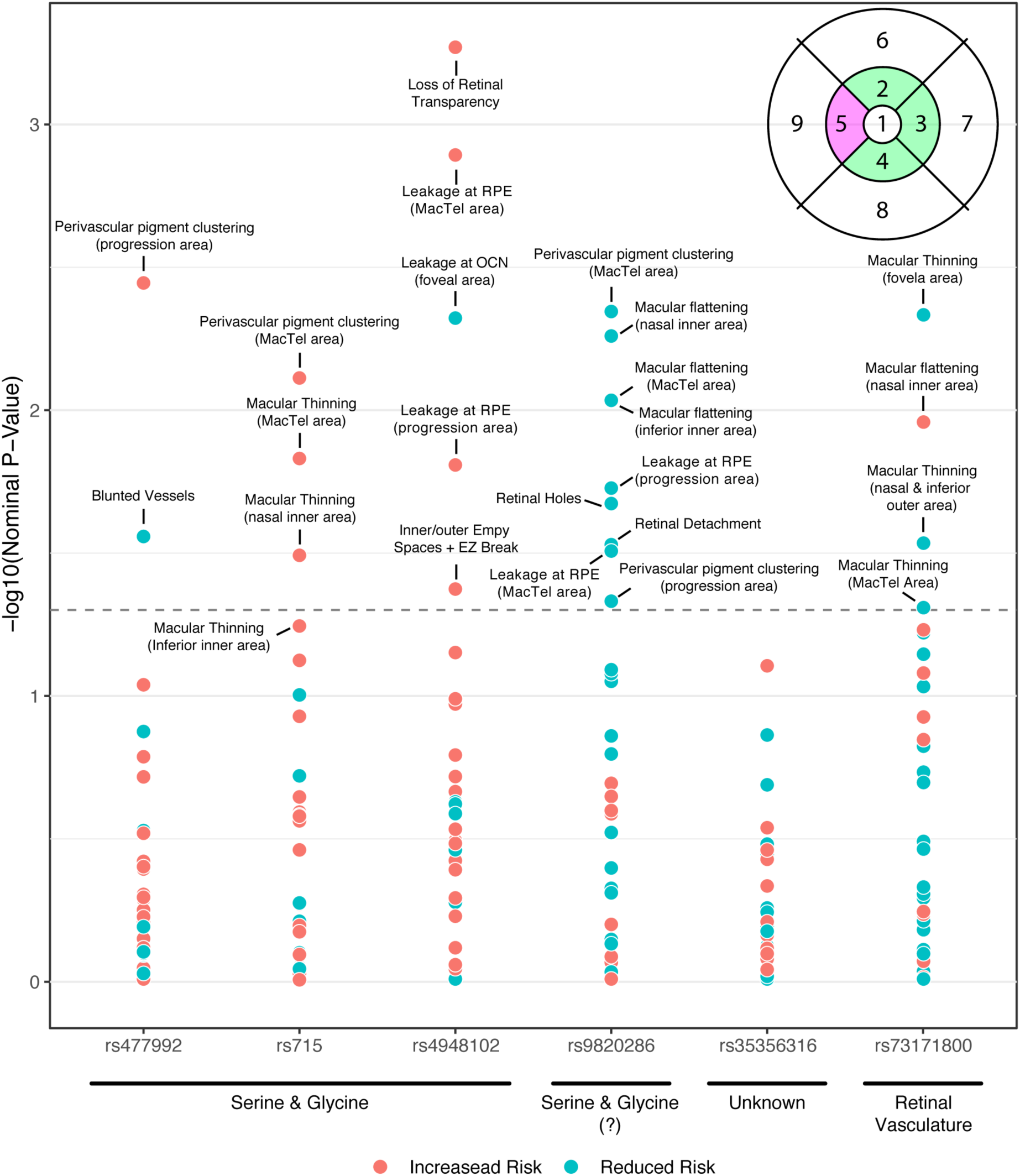
Association plot summarising all nominally significant association between MacTel SNPs and the endophenotypes (**Table S6**). SNPs are divided into categories based on MR results. Each dot represents the nominal p-value for each association between all SNPs and endophenotypes. Red dots indicate that MacTel risk alleles for that SNP also increases the risk of the retinal phenotype, while blue dots indicate a reduced risk. The dashed line represents the nominal significance of p=0.05. The top right corner displays the ETDRS grid for the right eye (OD) ^29^ used to divide the endophenotypes into 9 retinal areas; 1: foveal area, 2: superior inner area, 3: nasal inner area, 4: inferior inner area, 5: temporal inner area, 6: superior outer area, 7: nasal outer area, 8: inferior outer area, and 9: temporal outer area. Although definitions vary across the literature, for simplicity in this manuscript we define the pink area as the ‘MacTel [onset] area’ while the green areas are MacTel ‘progression areas’.

These three loci, which converge on glycine/serine metabolism (1p12/rs477992/*PHGDH*, 2q34/rs715/*CPS1*, and 7p11.2/rs4948102/*PSPH*) all demonstrated an overall susceptibility increment to RPE endophenotypes and other macular phenotype risks, lending further support for their role, despite not attaining significance after p-value adjustment.

Interestingly, the MacTel risk allele at locus 3q21.3 (rs9820286) was only associated with protection from retinal disease endophenotypes. Specifically, locus 3q21.3 was associated with protection against retinal holes, retinal detachment, perivascular pigment clustering in the MacTel area and progression areas; leakage at the RPE level in the MacTel area and progression area; and foveal slope flattening in MacTel area, inferior inner area, and nasal inner area.

Locus 5q14.3 (rs73171800) corresponded to protection against macular thinning in the foveal area and nasal-inferior outer areas and increased risk of foveal flattening in the nasal inner area. Clustering the loci using hierarchical clustering on their regression coefficients (**Figure S6**) highlighted the unique endophenotype effect profiles of the 5q14.3 (rs73171800) and 3q21.3 (rs9820286) loci, and their distinctiveness in comparison to the other three loci, known to act on the serine/glycine pathway.

## Discussion

This study exploited well-powered publicly available GWAS data from traits of interest as well as deep retinal phenotyping data to further investigate the genetic aetiology of MacTel, a rare retinal disorder for which the first GWAS was only recently performed^10^.

Mendelian randomization and publicly available data were used to “genetically predict” abundances of different metabolites. Although all GPMs in this study were generated using estimated SNP allele dosage effects from blood-based metabolomic studies rather than retina, we believe that our results represent a useful approach for interrogating other retinal disorders.

We found serine depletion as the strongest causal driver of MacTel risk, with an association magnitude much greater than those observed when using single SNPs. The direction of association between GP serine and MacTel agrees with results from direct serum serine measurements reported previously^10^, compelling additional evidence for a causal association between serine deficiency and MacTel. MacTel disease odds doubled for every standard unit decrease in GP serine, and individuals in the lowest quintile of GP serine presented an OR of 6.34 for MacTel compared to individuals in the top quintile. Our results also confirm the pronounced serine role in disease progression. As this analysis was applied only to MacTel patients and heterogeneity of retinal phenotypes may be under-represented, GP serine was nevertheless able to partly discriminate between subjects with more advanced retinal abnormalities and those without. For example, lower GP serine had a clear association with retinal greying in all retinal areas, a phenotype not observed in all MacTel patients^4^. GP serine also affected retinal thinning in the temporal parafoveal area, a marker of photoreceptor degeneration. Our results suggest a highly plausible biochemical explanation for differences in disease heterogeneity and progression. However, we acknowledge that our study used only aggregated retinal phenotypes and proxy measures of progression. Longitudinal data documenting progression is now required to confirm these findings. High concentrations of deoxy-sphingolipids, a byproduct of serine deficiency, have recently been shown to cause MacTel^15^. Our results provide evidence that genetically-encoded serine depletion is causal for MacTel, which likely contributes to the disease by promoting deoxy-sphingolipid biosynthesis^15^.

Our results indicate that the association between GP glycine and MacTel is likely an artefact of the shared genetic signal between this metabolite and serine. This may relate to the considerably greater sample size used to construct the glycine GP, and/or its metabolic dependence on serine. Similarly, the previously observed significant association of threonine to MacTel disease risk^10^ is likely due to its metabolic dependence on glycine and serine and thus a bystander effect.

MTAG confirmed the central role of locus 7p11.2 (rs4543497), which was previously found to be only nominally significant, and identified a new locus, 8q24.1 (rs2954021). rs2954021 is predictive of endogenous serum serine concentrations^22^ and thus might confer MacTel risk by contributing to the same serine pathway as previously identified SNPs. Conversely, we did not find any additional signal arising from shared genetic correlations with retinal vascular calibre traits.

Although not significant after correction for multiple testing, SNP rs73171800 affected macular thickness, specifically, a protective effect of the C allele against macular thinning, or an increase in macular thickness. Interestingly, a large GWAS study previously found that the G allele of SNP rs17421627 - corresponding to the C allele of SNP rs73171800 (LD with rs73171800 r^2^=0.67), was significantly associated with increased macular thickness^33^. A targeted study of SNP rs17421627 demonstrated enhancer activity which modifies the retinal vasculature^34^. Substituting the homologous zebrafish locus with a construct containing rs17421627 resulted in enhancer activity and changed expression of a proximal microRNA, mir-9-2 (homologous with human mir-9-5). Both mir-9-2 knock-down and endogenous enhancer knock-out animals showed dysmorphic retinal vasculature, indicating that rs17421627 may act on miRNA expression to modify the formation of the retinal vasculature in humans. Although we did not find a convincing causal relationship between retinal vascular calibre and MacTel it may be that other features of the vasculature, for example, leakage, branching or integrity, are modified in rs17421627 carriers, and account for the gross differences in macular thickness. Further deep phenotyping of the retinal vasculature in these individuals may yield the physiological basis for this phenotype and its impact on MacTel.

By conditioning on GP serine, glycine and T2D we revealed independent genetic contributors to MacTel. The locus 3p24.1, tagged by SNP rs35356316, lies between *EOMES* (<u>604615</u>) and *SLC4A4* genes (<u>603345</u>). Interestingly, *EOMES* encodes a transcriptional activator which is shown in mouse studies to interact with the retinal transcription factor *Pou4f2*^35^. The downstream gene *SLC4A4* encodes a sodium-coupled bicarbonate transporter which is expressed in Müller glia and the RPE, and functions to balance pH in the subretinal space^36^.

Conditional analysis revealed SNP rs36259 at locus 19p13.2 which tags an exonic, non-synonymous SNP located within the gene *CERS4* (<u>615334</u>), encoding a dihydroceramide synthase involved in sphingolipid biosynthesis^37^. Just as serine depletion is associated with defective sphingolipid synthesis, this locus may reduce sphingolipid production and thus contribute to MacTel.

Type 2 diabetics are overrepresented among MacTel patients^16^. We found a weak positive association between GP T2D and MacTel risk. A possible explanation for this is that T2D involves major perturbations of patient metabolism. Specifically, a recent meta-analysis of metabolite abundances associated with pre-diabetes and/or T2D, found that glycine depletion tends to occur in this disease^38^. We show that glycine depletion is not likely to be causative for MacTel and is likely to be a consequence of metabolomic co-regularisation counterbalancing the genetically induced serine depletion. We speculate that diabetes may be a consequence rather than a cause of the metabolic phenotype underlying MacTel. Future MR studies may further dissect this interaction but will require a more highly powered MacTel GWAS. Indeed, T2D risk remained significant even after controlling for GP serine and GP glycine, which indicates the possibility of a separate mechanism unrelated to glycine/serine metabolism.

Our analysis revealed that locus 3q21.3, previously believed to act on the disease by modifying serum glycine and serine, is instead likely disconnected from these metabolites, as is the case for loci 3p24.1 and 5q14.3, implying that additional potential disease mechanisms exist.

Future studies of disease progression and experimental validation of serine-independent contributions will help to resolve the complex aetiology of MacTel and hence target treatments leading to increased efficacy.

## Acknowledgments

We would like to acknowledge the funding support from the Lowy Medical Research Institute. This work was also made possible through the Victorian State Government Operational Infrastructure Support and Australian Government National Health and Medical Research Council (NHMRC) independent research Institute Infrastructure Support Scheme (IRIISS). RB was supported by the Melbourne International Research Scholarship. BREA was supported by an NHMRC early career Fellowship (1157776). MB was supported by an NHMRC Senior Research Fellowship (1102971) and Program Grant (1054618). We are grateful to Dr Luca Lotta and Dr Claudia Langenberg for sharing their summary results metabolomics data and their extremely helpful contributions. Dr Saskia Freytag, Dr Anna Quaglieri, Prof Terry Speed, Prof Gordon Smyth, Dr Mari Gantner, Dr Kevin Eade, Dr Martina Wallace, A/Prof Christian Metallo, Prof Martin Friedlander, Dr Tjebo Heeren, Dr Mali Okada, Dr Sasha Woods, Prof Marcus Fruttiger, and Dr Catherine Egan for their extremely helpful contributions that greatly improved the quality of this manuscript. We are especially grateful to Dr Xueling Sim and Prof Wong Tien Yin for kindly sharing their summary data on retinal vascular calibre traits.

## Supplementary Material

### Supplementary Images

**Figure S1:**
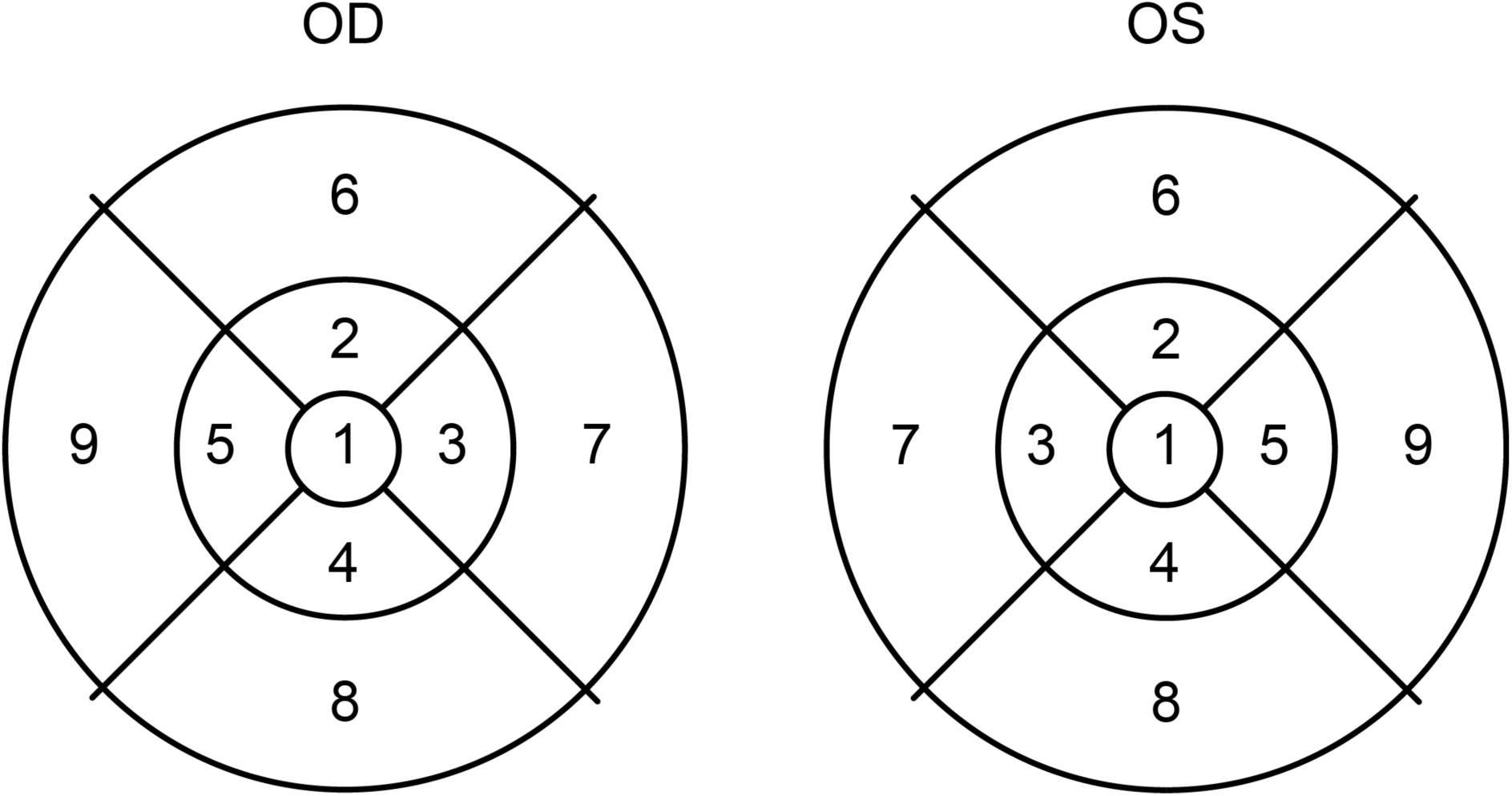
Right (OD) and left (OS) eyes macula, were divided into 9 areas according to the ETDRS grid ^1^. 1: foveal area, 2: superior inner area (progression area), 3: nasal inner area (progression area), 4: inferior inner area (progression area), 5: temporal inner area (MacTel area), 6: superior outer area, 7: nasal outer area, 8: inferior outer area, and 9: temporal outer area.

**Figure S2:**
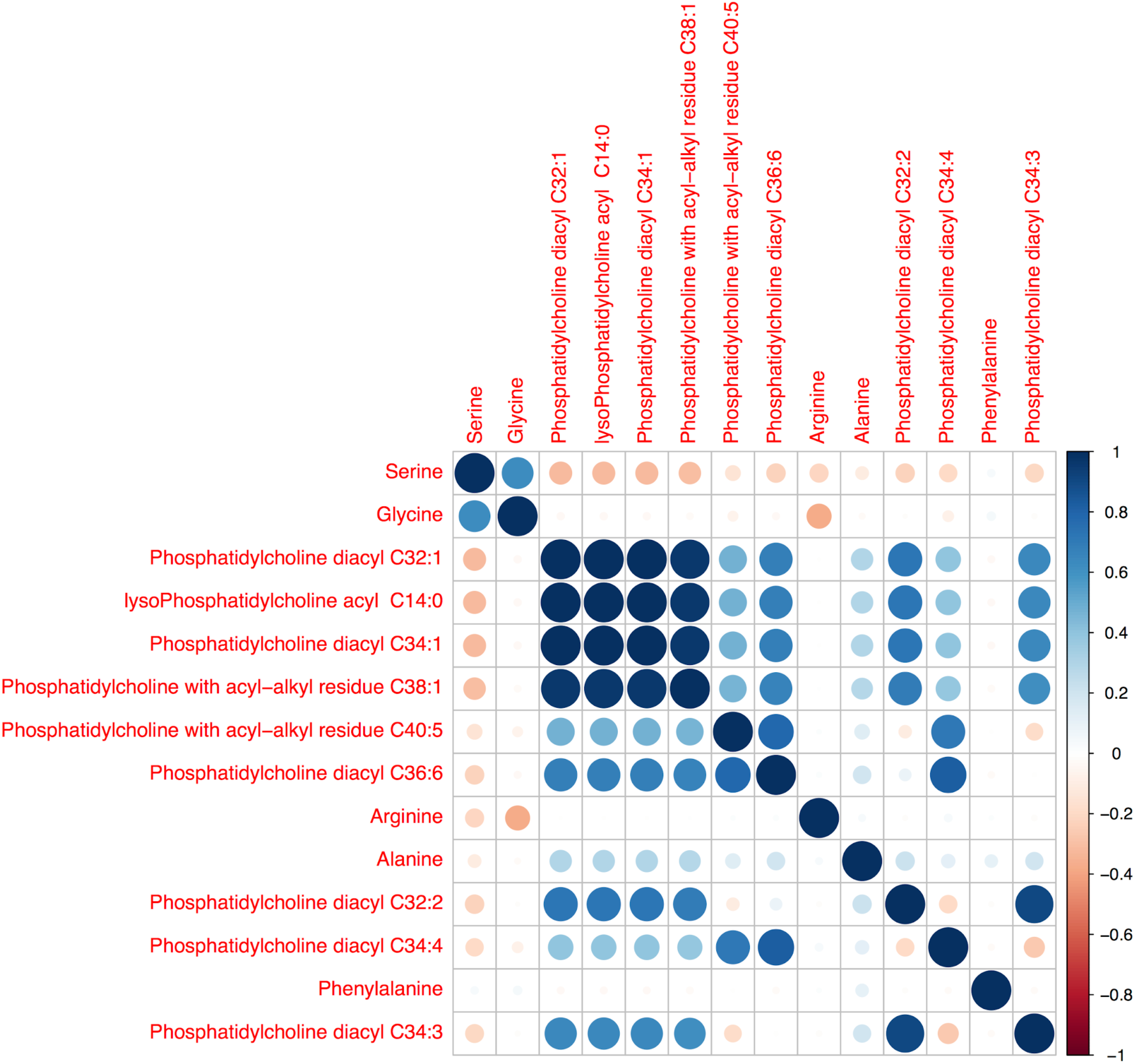
Correlation between GPM scores for all GPM scores that achieved multiple testing corrected significance (p < 0.05). Bigger dots and saturated colour indicate stronger correlations. Blue indicates a positive correlation while red indicates a negative one.

**Figure S3:**
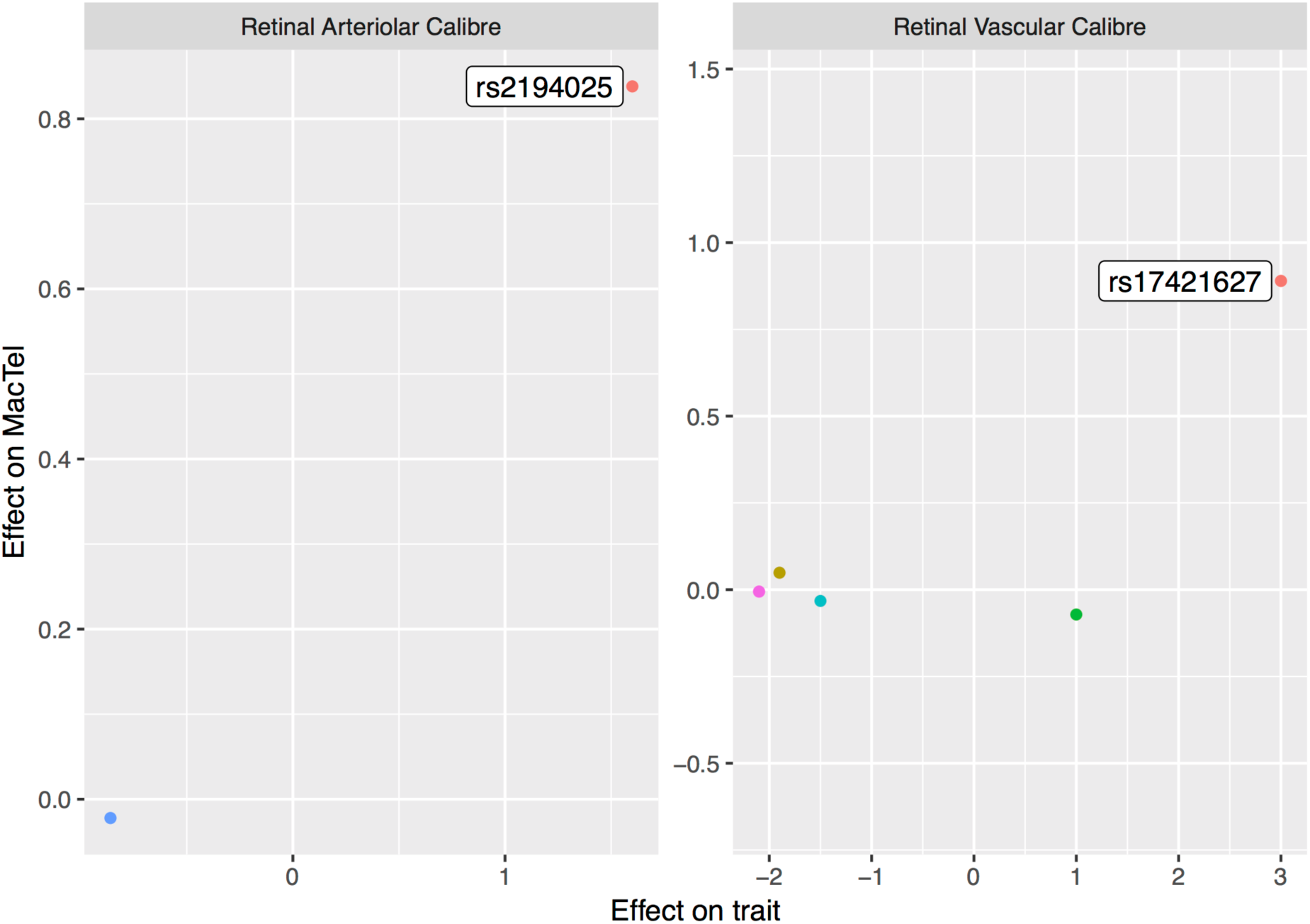
Comparison between SNPs effect on vasculature traits and MacTel. Each dot is a SNP which was found to be significantly associated with a vasculature trait. The x-axis represents the effect that the SNP has on the particular trait while the y-axis displays the effect that the SNP has on MacTel. Most SNPs have no effect on MacTel. The only two SNPs that have an effect are the SNPs rs2194025 and rs17421627 which are very close to each other and in strong LD with SNP rs73171800 (r^2^=0.94 and r^2^=0.67).

**Figure S4:**
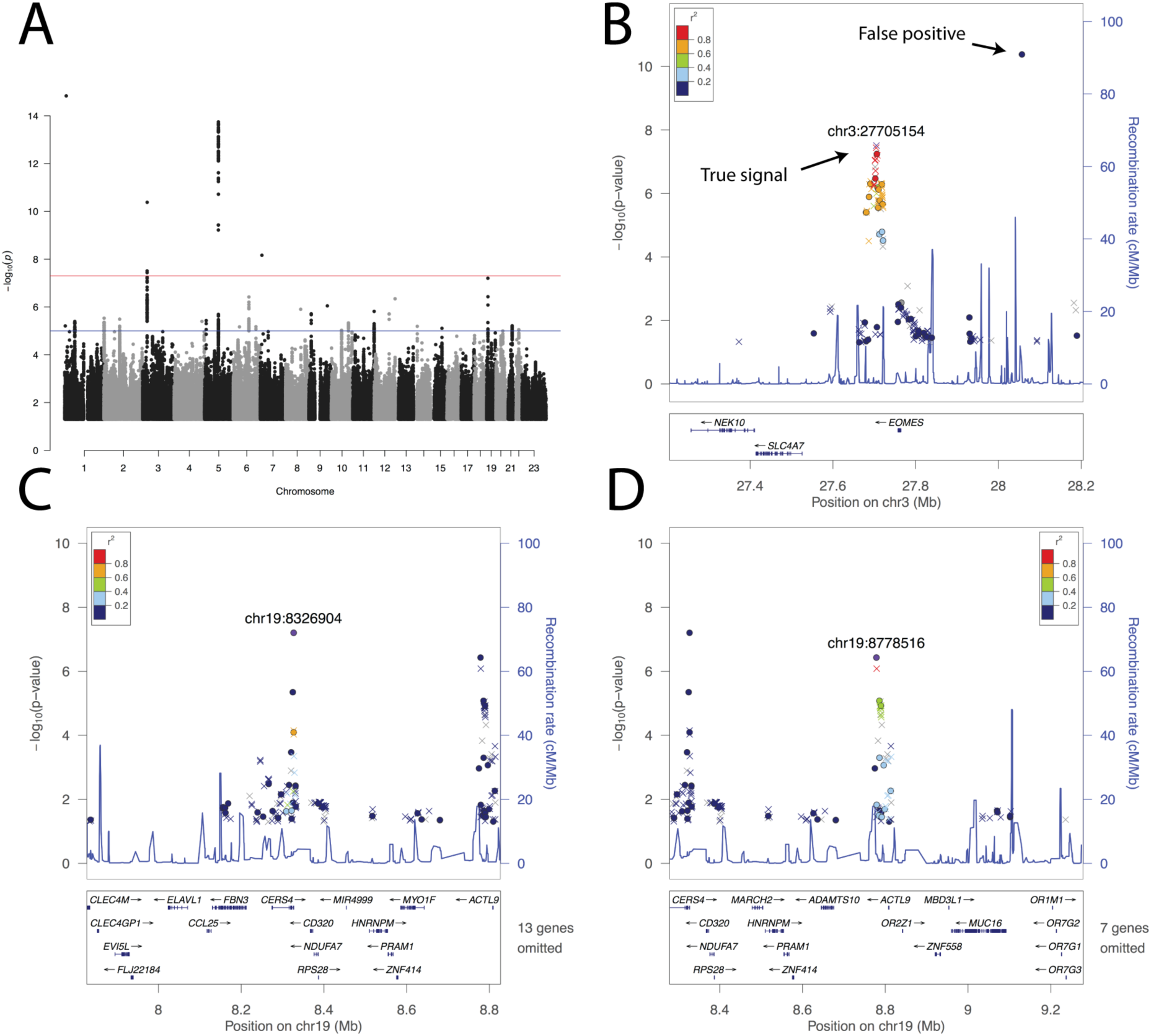
MacTel GWAS results conditioned by GP serine and GP glycine. A) Manhattan plot. B) Locus zoom plot of the significant signal on locus 3p24.1. C) Locus zoom plot of the significant signal by SNP rs36259 on locus 19p13.2. D) Locus zoom plot of the significant signal by SNP rs4804075 on locus 19p13.2.

**Figure S5:**
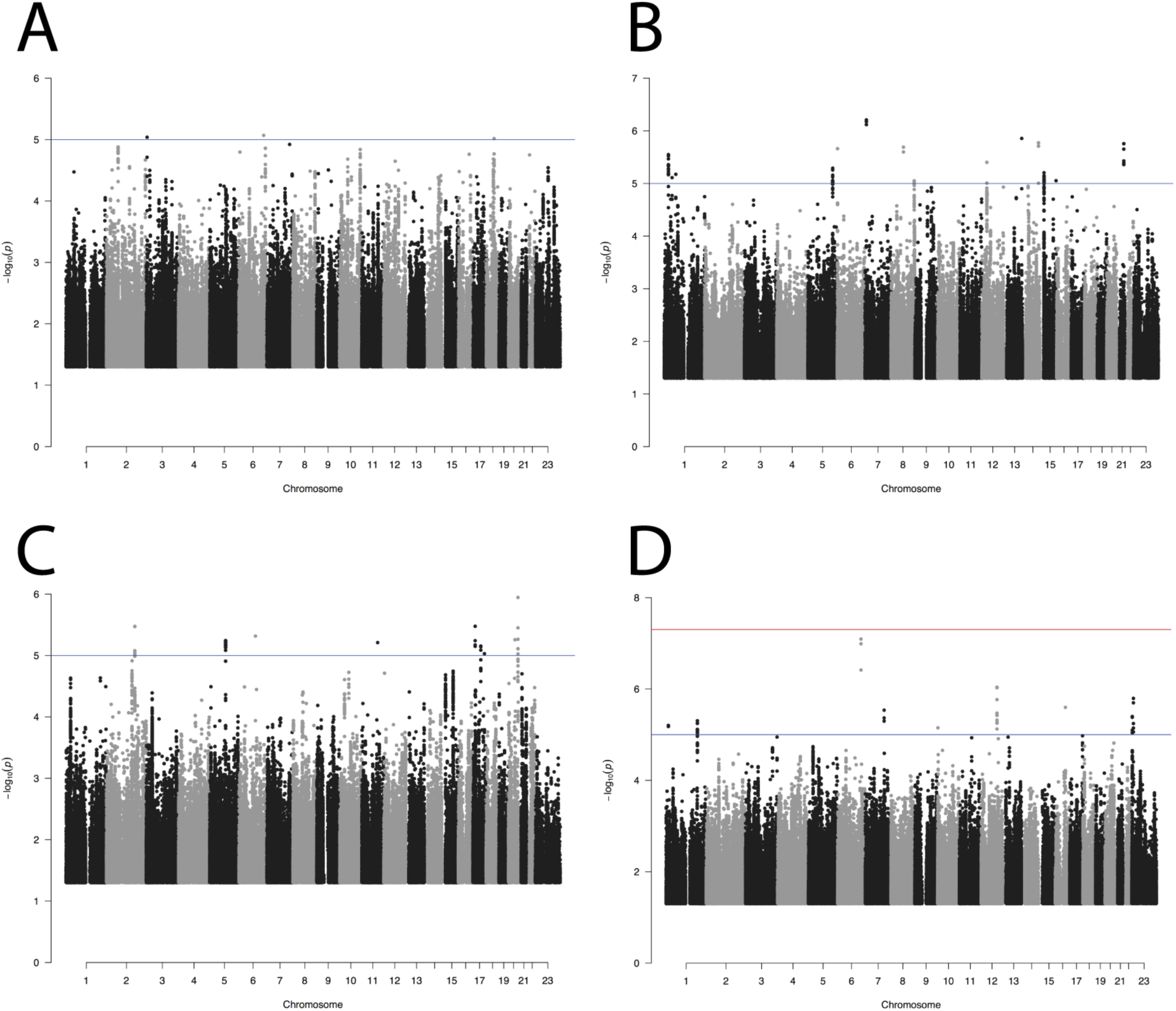
MacTel interaction GWAS results. A) Manhattan plot showing SNP interactions with GP serine, B) Manhattan plot showing SNP interactions with GP glycine, C) Manhattan plot showing SNPs interaction with SNP rs73171800 in locus 5q14.3, D) Manhattan plot showing SNP interactions with GP T2D.

**Figure S6:**
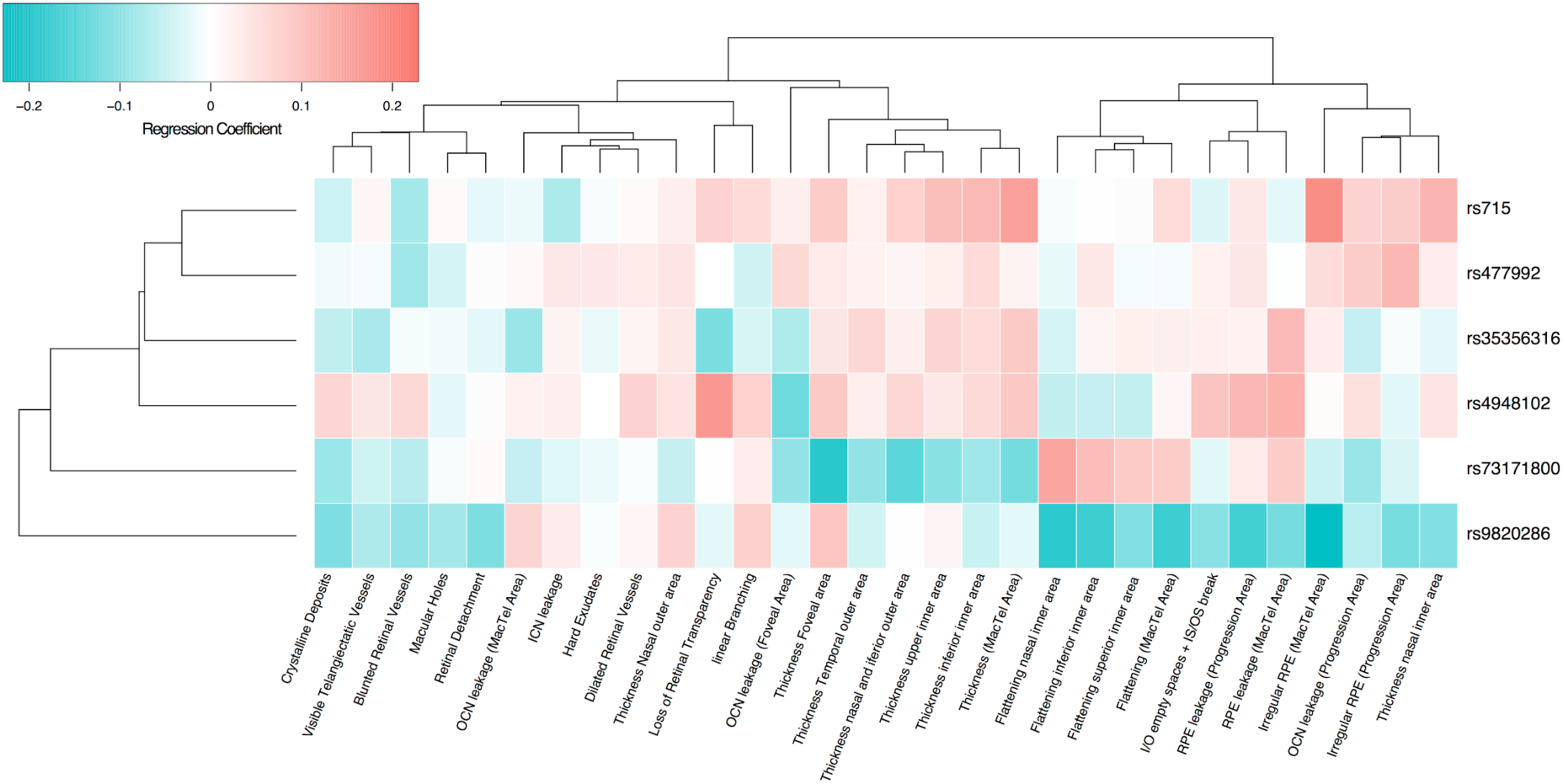
Loci clustering heatmap. This clustering heatmap visualises the regression coefficients of each SNP on the endophenotypes. Rows and columns of the heatmaps were ordered according to the hierarchical clustering to give a visual representation of the similarities. SNP rs73171800 (5q14.3) and SNP rs9820286 (3q21.3) have very different effects from the other four SNPs which show a more consistent, shared pattern.

### Supplementary Tables

**Table S1:** SNPs and weights used to create GPMs, GPTs, Shin et al GPMs and MTAG serine (available as .xlsx file).

**Table S2:** List of all phenotypes and measurement methods used to create the endophenotypes (available as .xlsx file).

**Table S3:** List of all GPMs associations and conditional GPMs association results (available as .xlsx file).

**Table S4:** List of all Shin et al GPMs associations results (available as .xlsx file).

**Table S5:** Detailed FUMA output from MTAG analysis (available as .xlsx file).

**Table S6:** Endophenotypes analysis results (available as .xlsx file).

**Table S7:** Endophenotypes compositions and weights (available as .xlsx file).

**Table S8:** Members of the MacTel Project consortium (available at the end of this file).

## Supplementary Methods

### GPM score calculation

We used the formula *GP*(*M*/*T*)*_*i*_* = ∑*_j_*_∈*i*_ *m_j_* ∗ *SNP_j_*, where *GP*(*M*/*T*)_*i*_ is the genetically predicted metabolite (*M*) or trait, *Ti*. *SNP_j_* is the *j^th^* SNP that was significantly associated with the metabolite or trait *i*. *m_j_* is the magnitude (weight) that the *SNP_j_* has on the metabolite or trait. The weight *m_j_* was obtained as *m_j_* = *β_j_* where *β_j_* is the linear coefficient of the regression of *SNP_j_* on the metabolite or trait *i*. We used a total of 499 SNPs to predict the abundances of 142 metabolites in blood serum, 37 SNPs to construct a genetically predicted T2D risk, 5 for retinal venular calibre and 2 for the retinal arteriolar calibre. GPMs and GPTs were normalized to have a mean of zero and unit standard deviation.

### Imputation of phenotypic data

The phenotyping dataset contained multiple missing data points. Most missingness was related to imperfections in the ophthalmological images. By assuming missingness at random we imputed the missing data using predictive mean matching through chained equations, available in the package MICE ^2^.

### Generation of Endophenotypes with Factorial Analysis

The phenotypic data was summarised into endophenotypes. Endophenotypes have been defined as “measurable components unseen by the unaided eye along the pathway between disease and distal genotype”^3^. To identify and construct these underlying endophenotypes while ensuring that each only contained phenotypes strongly related to each other, we used an iterative factorial analysis algorithm. Given the very different nature of the phenotypes observed we performed two separate analyses for the creation of the endophenotypes one to capture macular thickness endophenotypes, and the other to capture macular abnormality phenotypes.

Starting with 43 phenotypic measurements for macular thickness and 76 for macular abnormality phenotypes we reduced these two data sets further, by only retaining correlated phenotypes within each dataset.

We discarded all those phenotypes/variables which:

- Had an absolute value of Spearman correlation with all other variables less than 0.3.
- Had an absolute value of Spearman correlation with the overall dataset score of less than 0.3.
- Would decrease the Cronbach’s alpha of the dataset

This procedure retained 33 phenotypes for macular thickness and 52 phenotypes for macular abnormalities. Variables that were retained after this step underwent an iterative factorial analysis procedure:

1. Perform a *parallel analysis* on the dataset to discover the number of factors *F*needed to summarise the phenotypic data
2. Run a *factorial analysis* with *F*number of factors
3. Rotate the factors using the *oblimin* rotation to allow for correlation between endophenotypes and obtain a loading value for each phenotype in each factor.
4. Create a sequence of loading thresholds *T* from 0.2 to 0.5 by 0.01
5. For each loading threshold*t* ∈ *T*and for each factor *f* ∈ *F*, create a dataset that contains all the phenotype that have a loading greater than *t*. This results in F datasets.
6. Calculate the Cronbach’s alpha in each dataset assessing the reliability of each factor.
7. Calculate the average of all factor’s alphas
8. Select the loading threshold *t* ∗ that maximize the average alpha.
9. Eliminate from the dataset all the variables that do not have any loading greater than *t* ∗.
10. Eliminate all variables which create factors by themselves.
11. Repeat 1-10 the entire process until no variables are removed.

This procedure was followed by an additional step, where we took the phenotypes that were discarded from the previous step and re-introduced into the endophenotype creation process, by themselves. Those phenotypes that were discarded a second time, were manually collapsed into meaningful new endophenotypes from previous clinical knowledge. These endophenotypes were obtained by simply summing the phenotypes values together.

Each factor created by this process was considered an endophenotype in further analysis. All endophenotypes resulted in continuous variables for which higher values represented negative impacts on patient’s eye health. Specifically, higher values of macular thickness endophenotypes represented a risk of macular thinning, higher values for foveal slopes endophenotypes represented increase risk of foveal flattening, and higher values for macular abnormalities endophenotypes represented an increased risk of observing that particular endophenotype. Most of the analyses to obtain the endophenotypes were performed using the R package Psych^4^.

Following the method used by ^5^, we also created foveal slope measurements from each parafoveal area using the macular thickness endophenotypes. The formula used is was *Slope*_*j*_ = *tan*^−1^((*Thick*_*j*_ − *Thick*_1_)/500). Where *Slope*_*j*_ indicates the slope from the fovea to the parafoveal area *j*and*Thick*_*j*_ is the endophenotype of macular thickness in area *j*. Since the foveal slope is defined as the slope from the fovea to the parafoveal areas the only areas used were area 2, area 3, area 4, and area 5 of the ETDRS grid.

The factorial analysis results in 9 endophenotypes describing macular thickness and 19 describing macular abnormalities. We discarded one macular thickness endophenotype and one macular abnormalities endophenotype since they were believed to describe only a residual correlation between phenotypes and have no real clinical relevance. Additionally, we obtained 4 endophenotypes describing macular slope leading to a total of 30 endophenotypes to be used in the analysis.

### Testing the identified disease key drivers and SNP loci against the endophenotypes

Endophenotypes were normalized to have zero average and unit standard deviation. Given the longitudinal nature of the data, we chose a linear mixed model approach to take into account the relationships within the subject and eye observation. Each model included all factors of interest as well as time defined as the number of days from the initial visit date, a quadratic function of visit year, gender, and age as covariates. Each model also contained a random intercept and random time slope for each subject as well as a random intercept and random time slope for each eye within each subject. We included all these random effects since the log-likelihood of models that contained them and models which did not were significantly different (results not shown. Models that compared fixed effects were estimated using restricted maximum likelihood estimation. Given the large sample size, the p-values were obtained by assuming the asymptotic normality property of the t-values for each association. To take into account false discovery rate we used an adaptive Benjamini-Hochberg procedure ^6^. We adopted this procedure rather than strict Bonferroni correction because of the very conservative nature of the linear mixed model and the small magnitude of SNP effect sizes.

## Supplementary Results

### Using MTAG to increase discovery power on GP serine

The much higher powered discovery GWAS for glycine and serine were used to predict glycine and serine GPMs. The samples sizes for these two traits were different: (nGlycine = 79,293 and nSerine=30,68). For this reason, we tried to increase the discovery power of the serine GWAS by using MTAG and combining serine GWAS results with the glycine GWAS results, leveraging the genetic correlation between them (**Figure 1A**). Although MTAG estimated the genetic correlation between the two traits to be high (0.413), the maximal FDR for serine was also high (0.246). This was most likely caused by the difference in sample sizes of the two studies. By using the results from this analysis, we constructed GP MTAG serine and tested it for significance alongside GP glycine on MacTel. This resulted in a slight decrease of significance for GP MTAG serine (p=6.24e-13) and a complete loss in significance for the glycine original GPM (p=0.907).

### MR using Shin et al 2014 metabolomics study

We also tried to perform MR testing on all metabolites available from one of the first studies on the genetic impact on metabolomics abundances by Shin et al 2014 ^7, 8^ which contained many more metabolites than those initially available to us. Of the 248 GPMs available we found few additional metabolites to be significant (**Table S4**), None of them was as strongly associated as GP glycine and GP serine and none retained significance after their inclusion as covariates.

### Composition of the endophenotypes

A detailed list of phenotypes and respective loadings values used to estimate each of the endophenotypes is available in **Supplementary Materials** and **Table S7**. Interestingly, among the 30 retained endophenotypes, we noticed that some distinguished between abnormalities occurring in the initial MacTel region (area 5 in the ETDRS grid) and abnormalities expanding to the surrounding areas of the retina, which is usually denoted to indicate progression of the disease (areas 2, 3 and 4 on the ETDRS grid). For this reason, we defined these endophenotypes as “MacTel area” and “Progression” respectively.

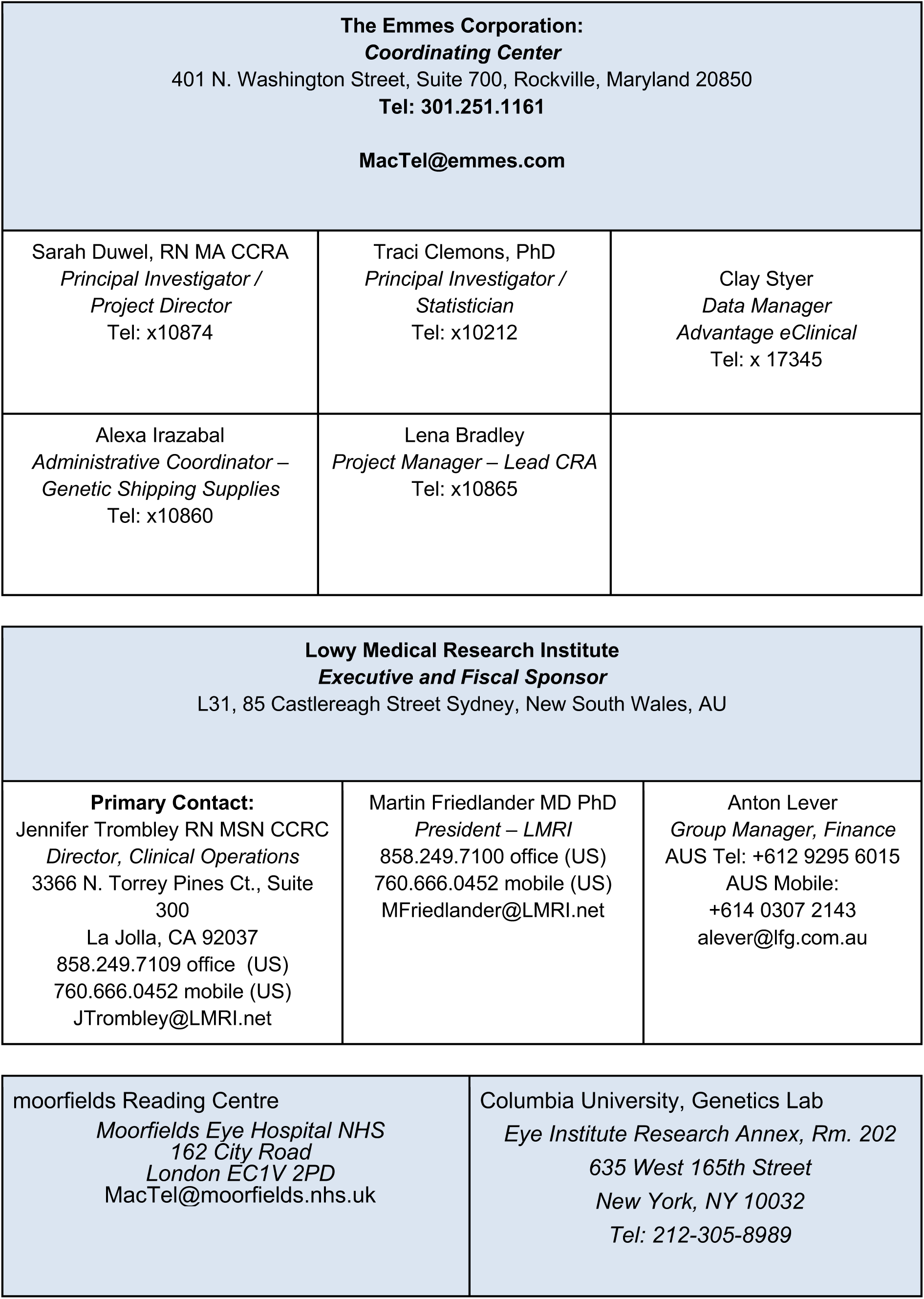
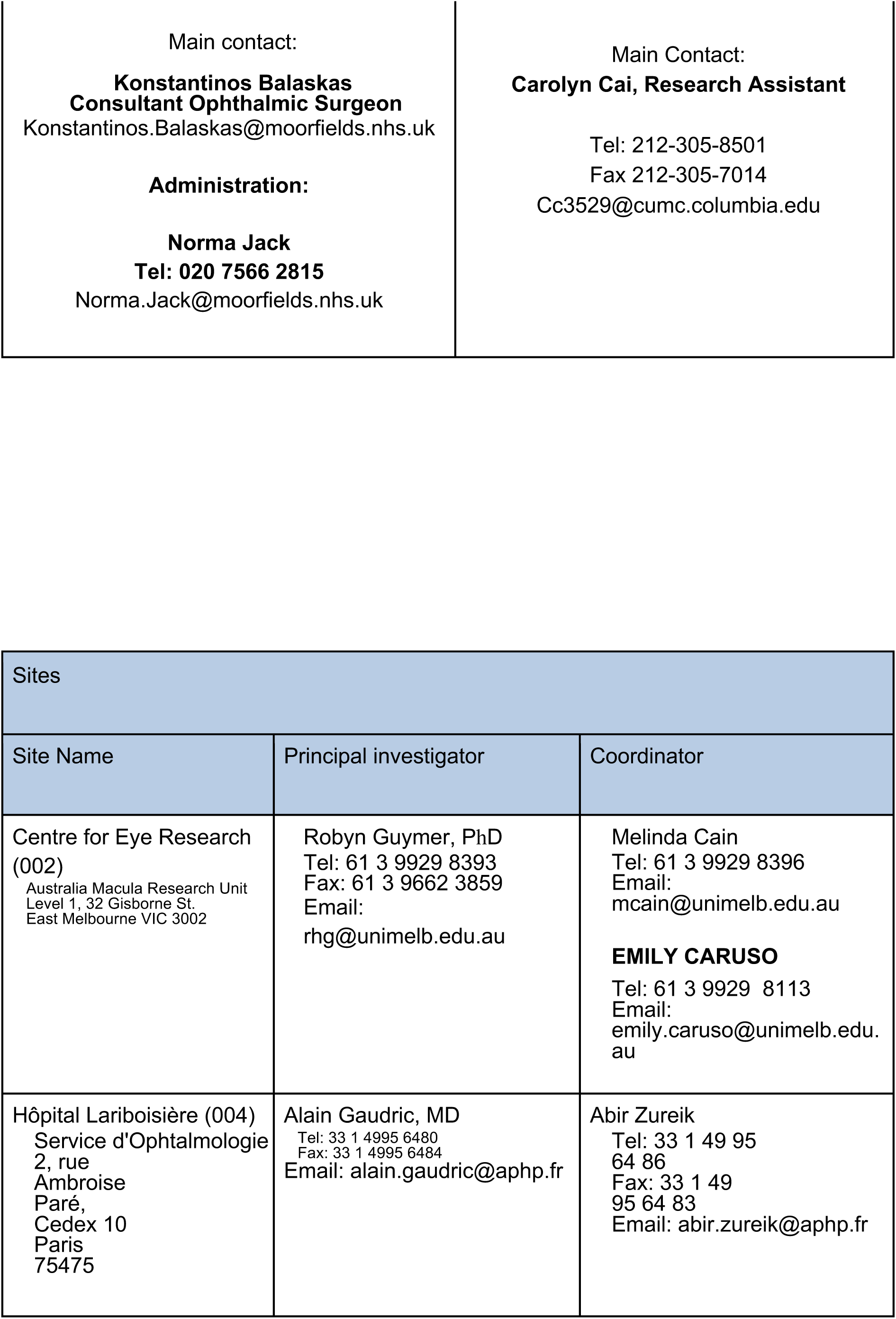
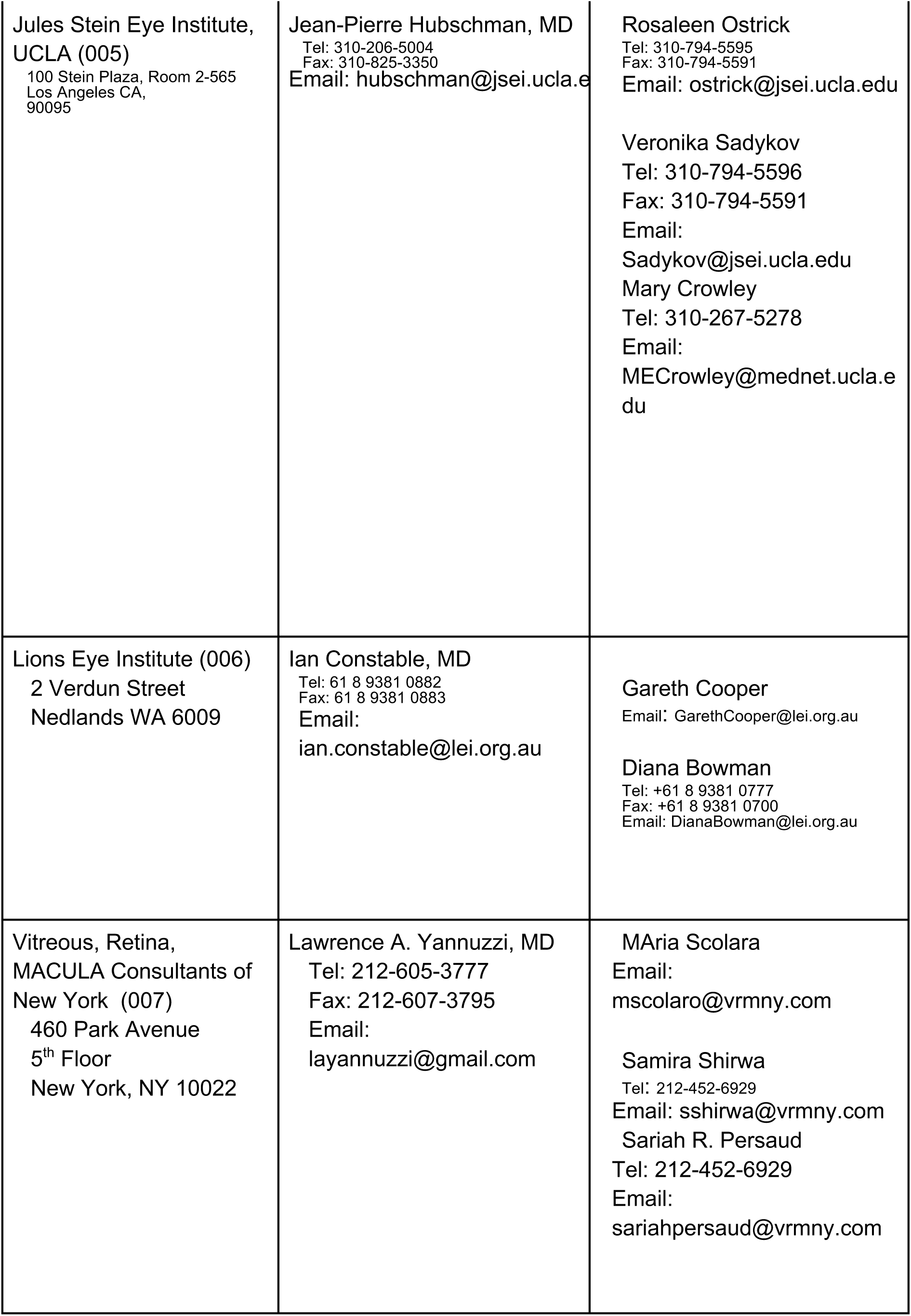
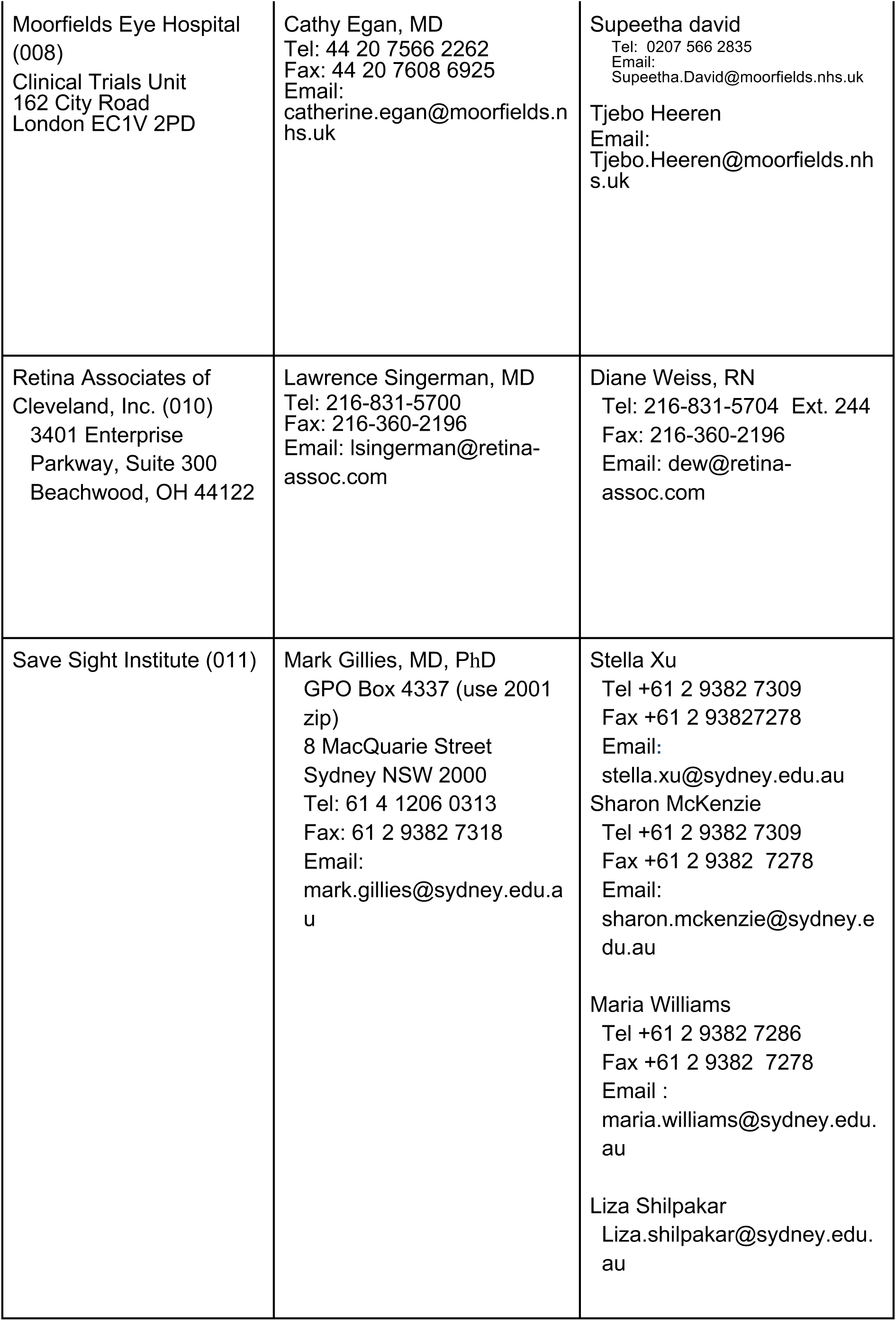
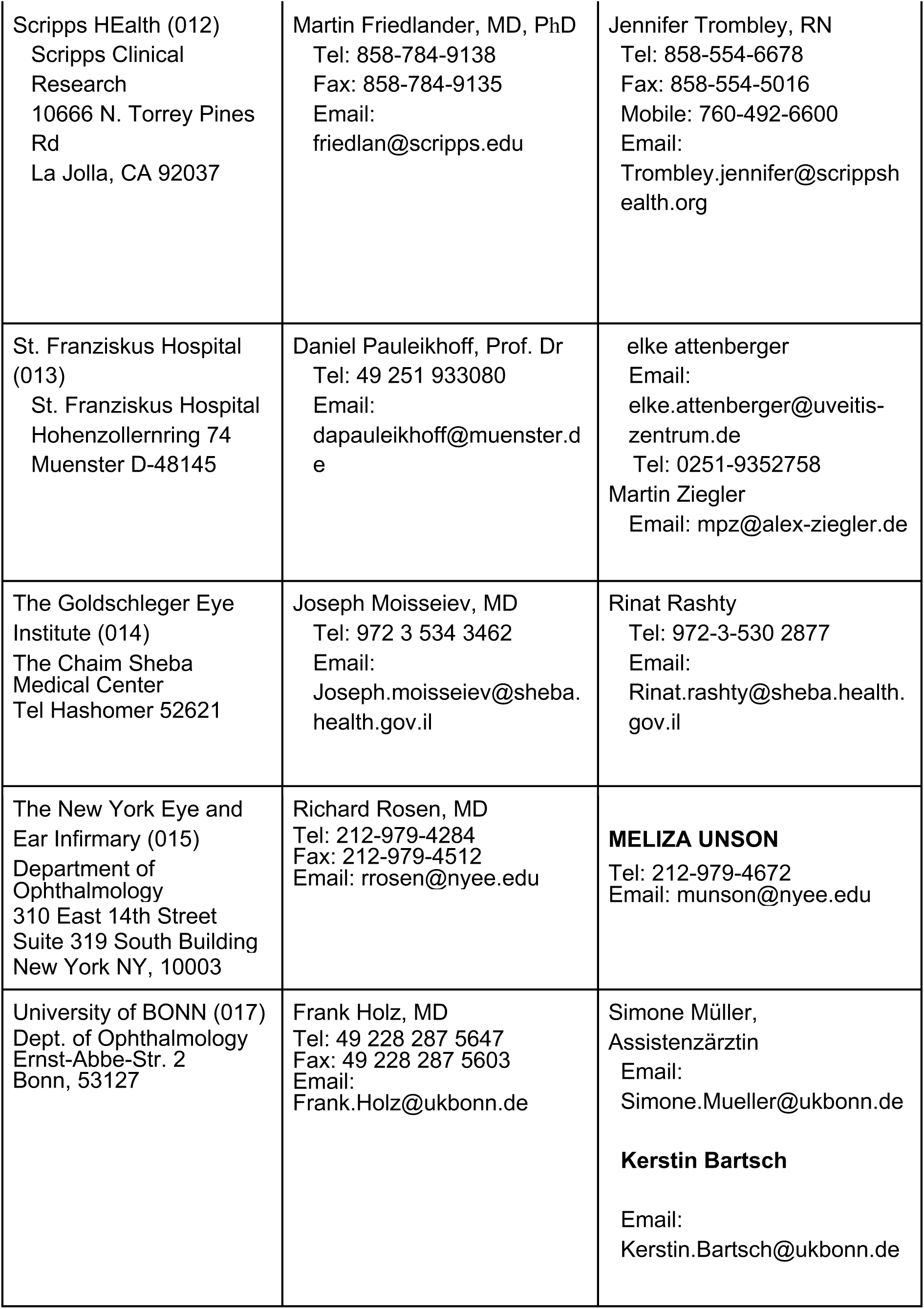
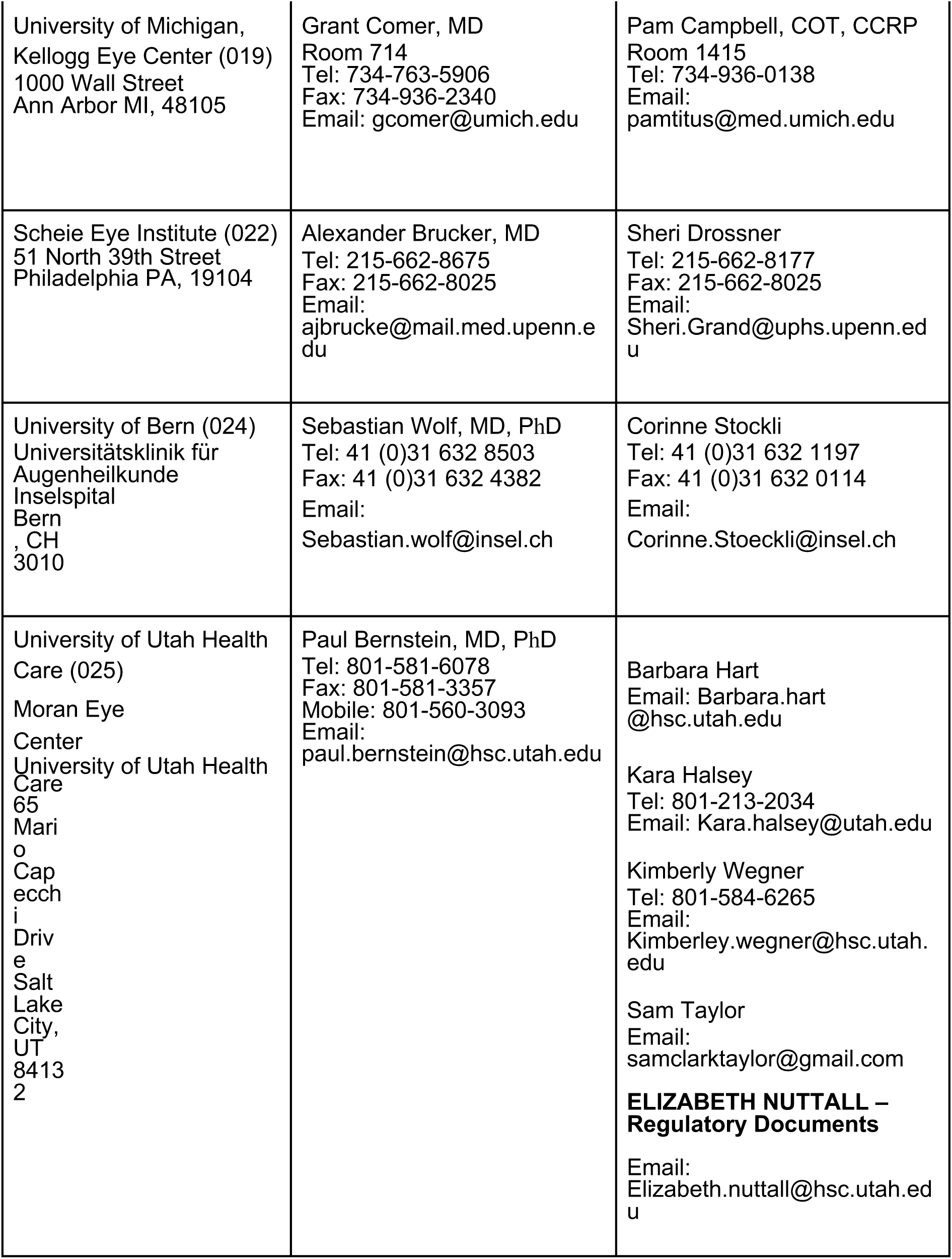
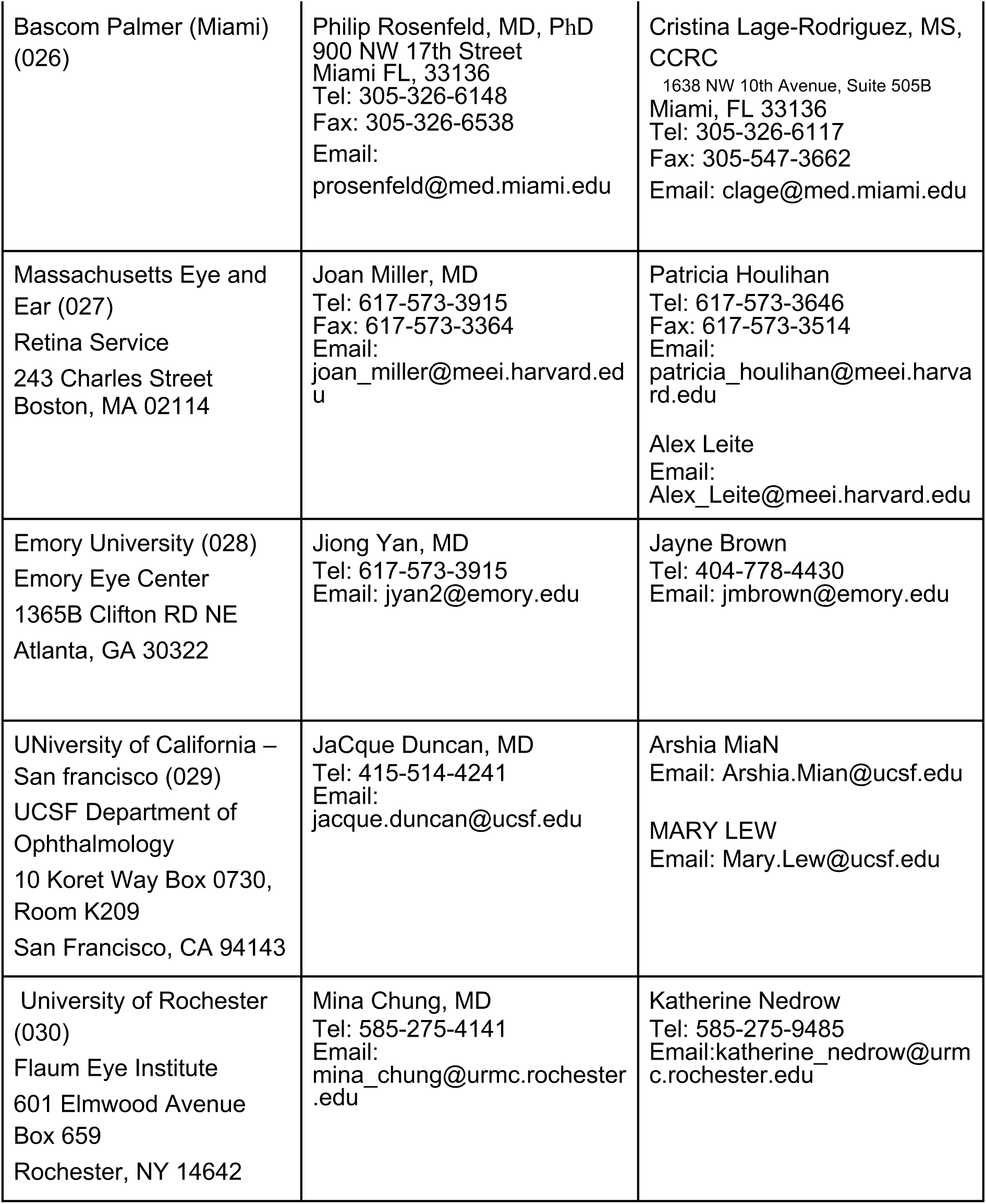
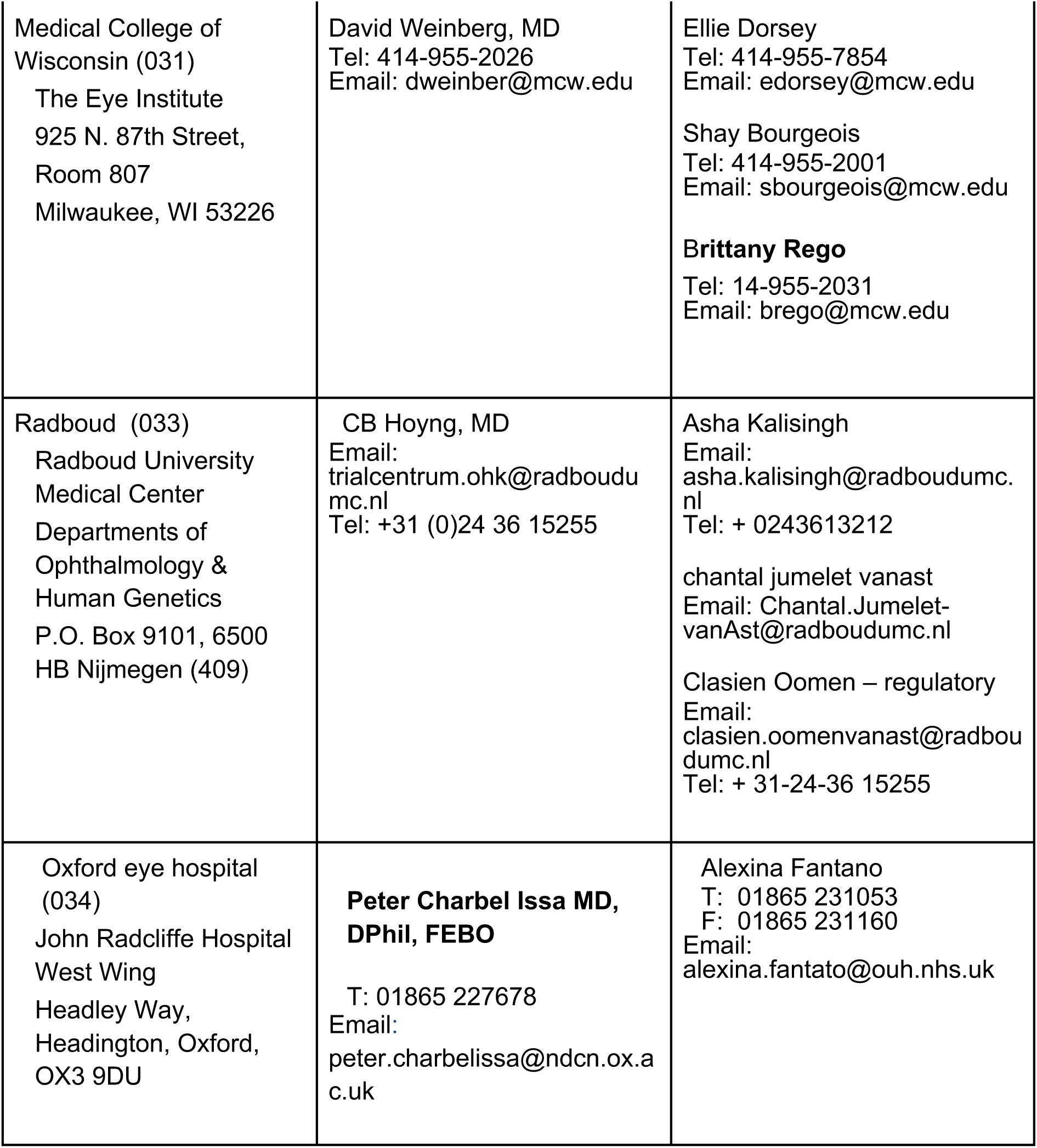
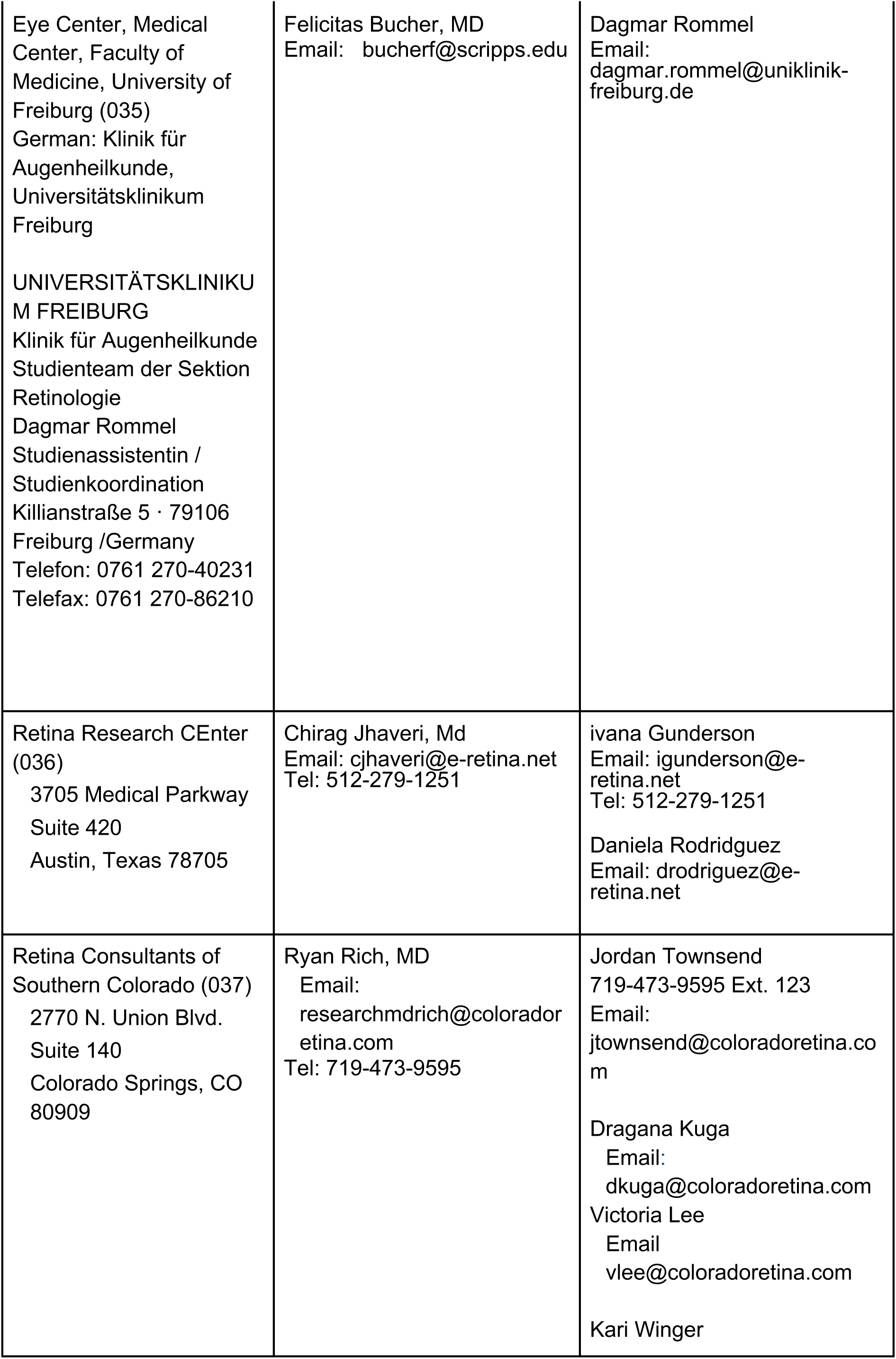
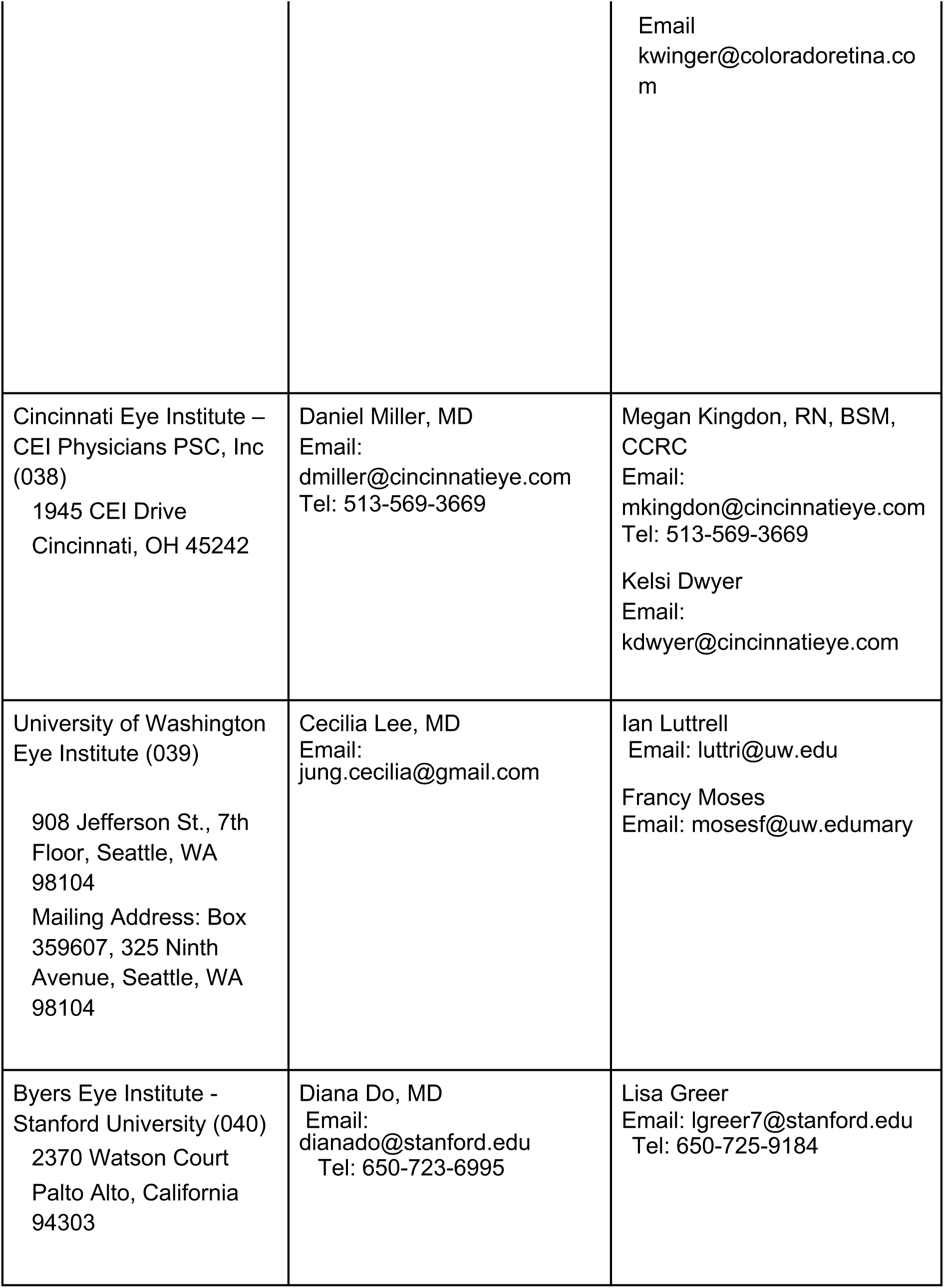
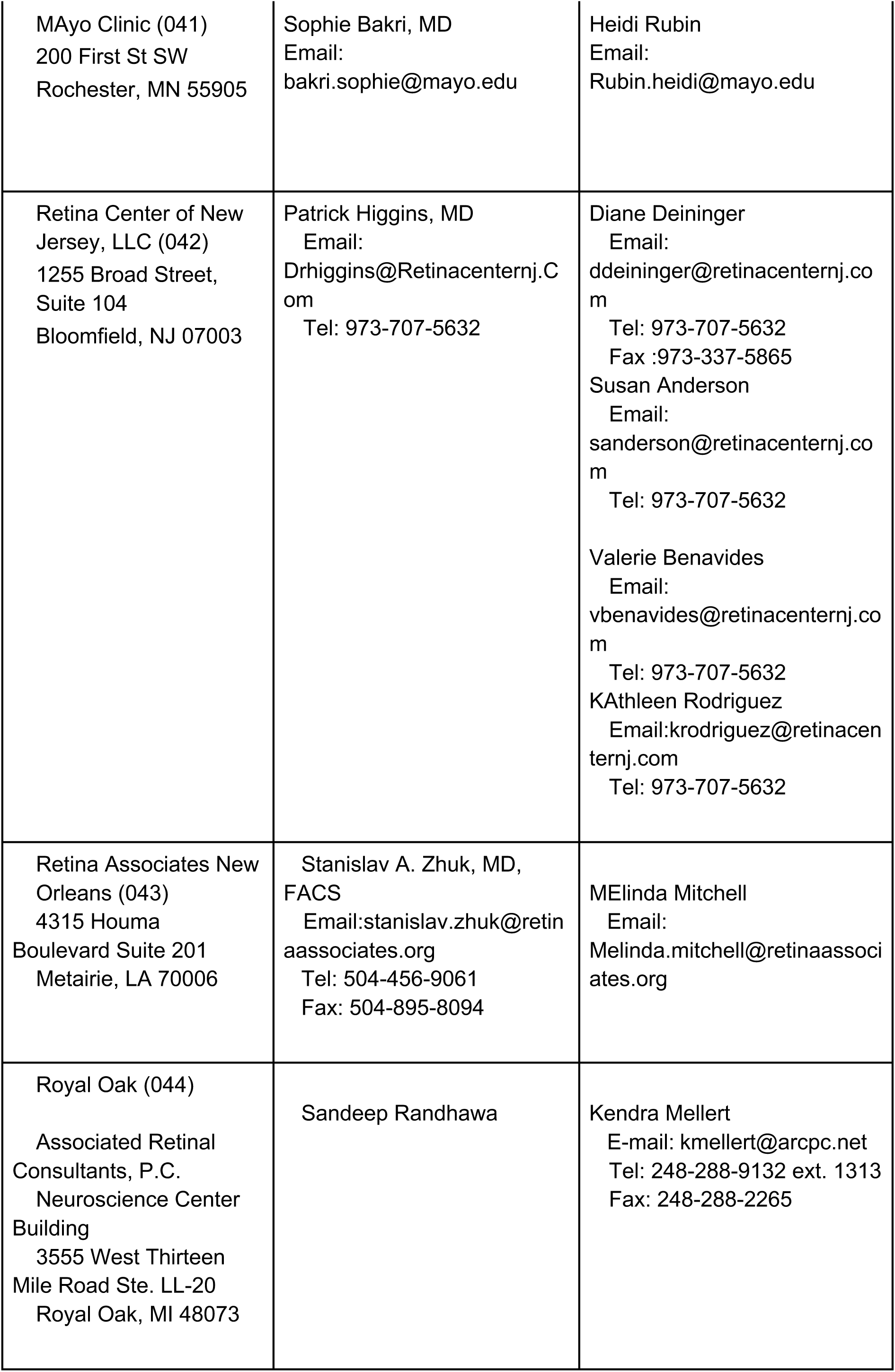
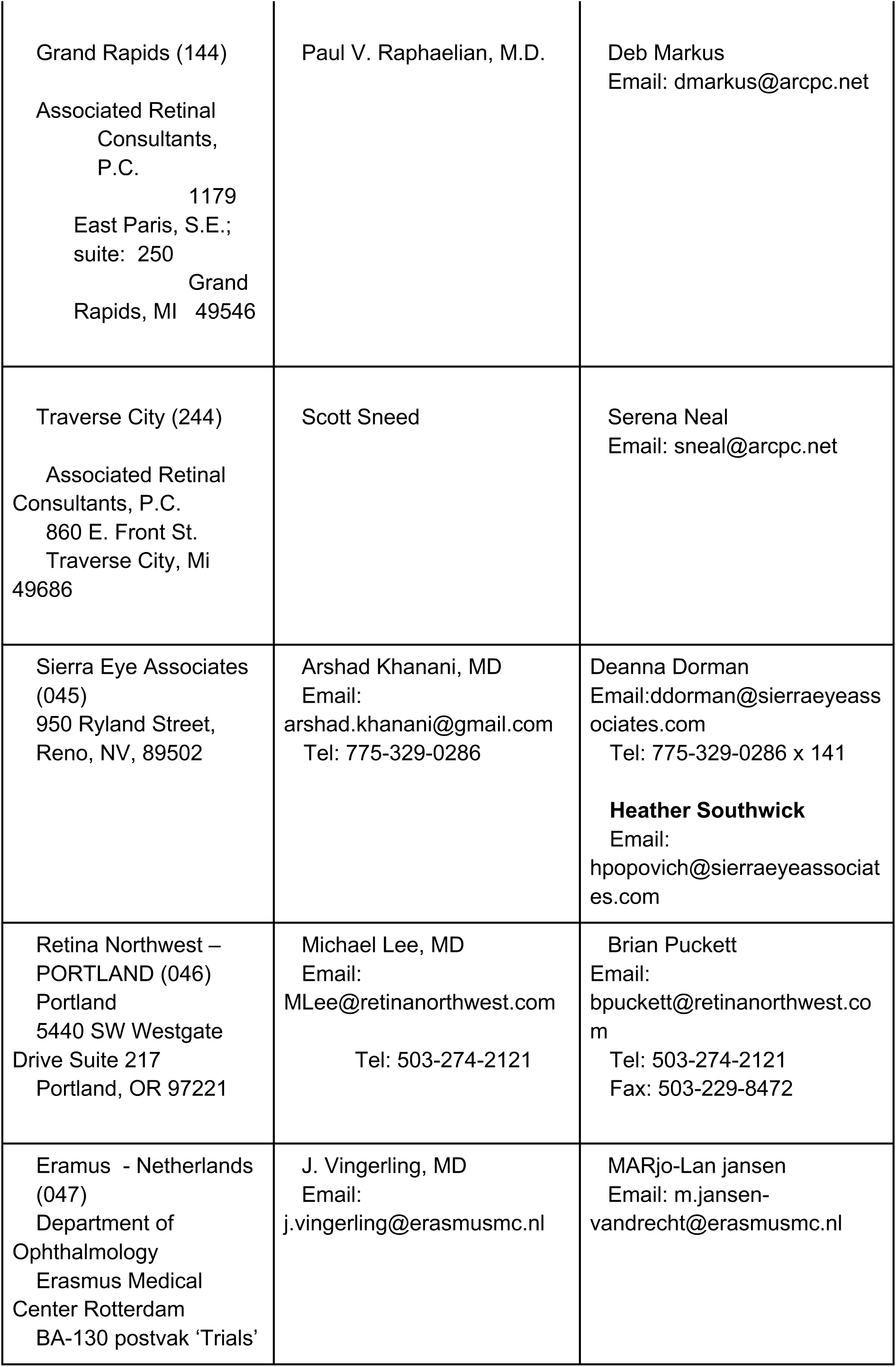
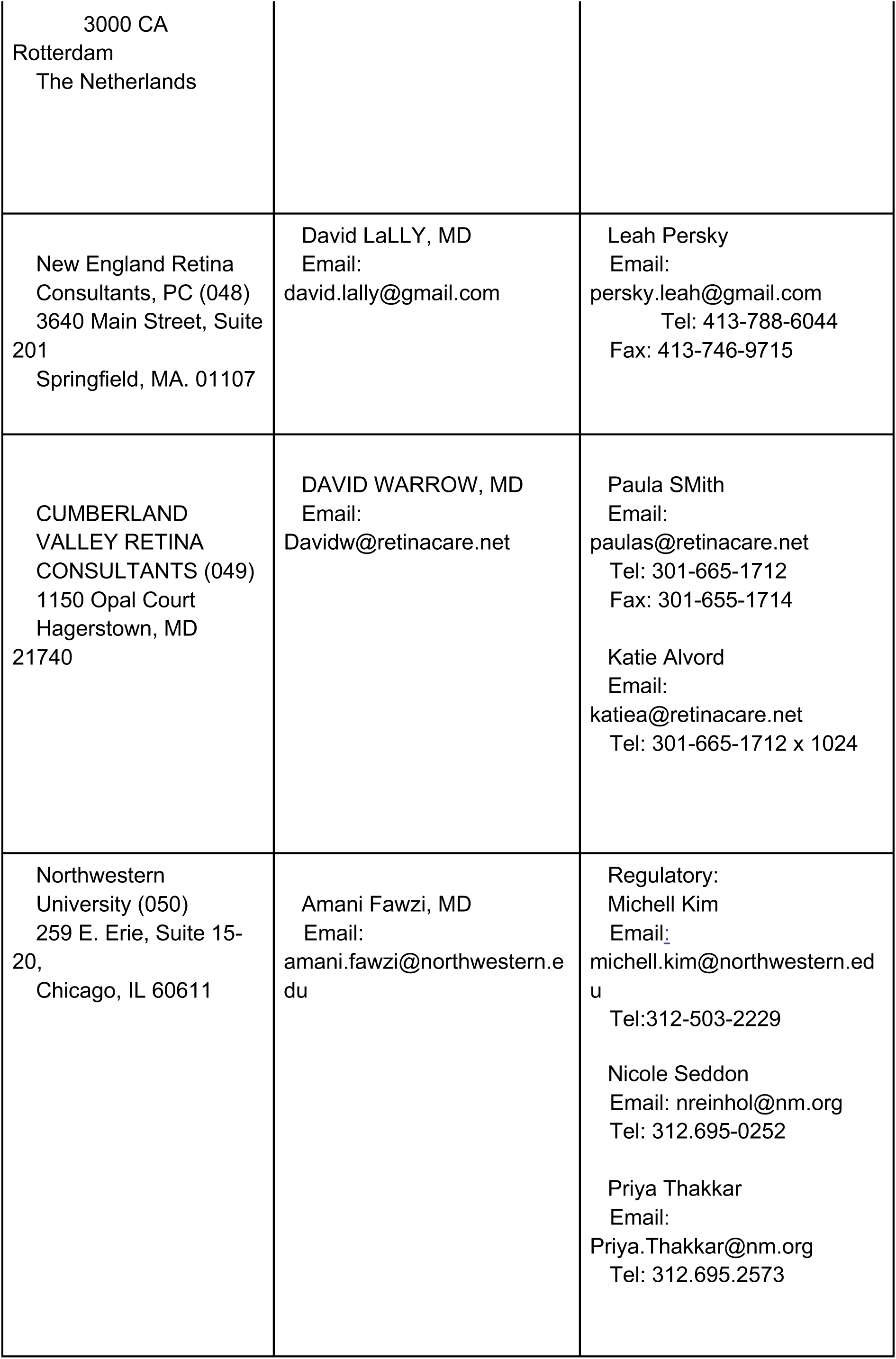
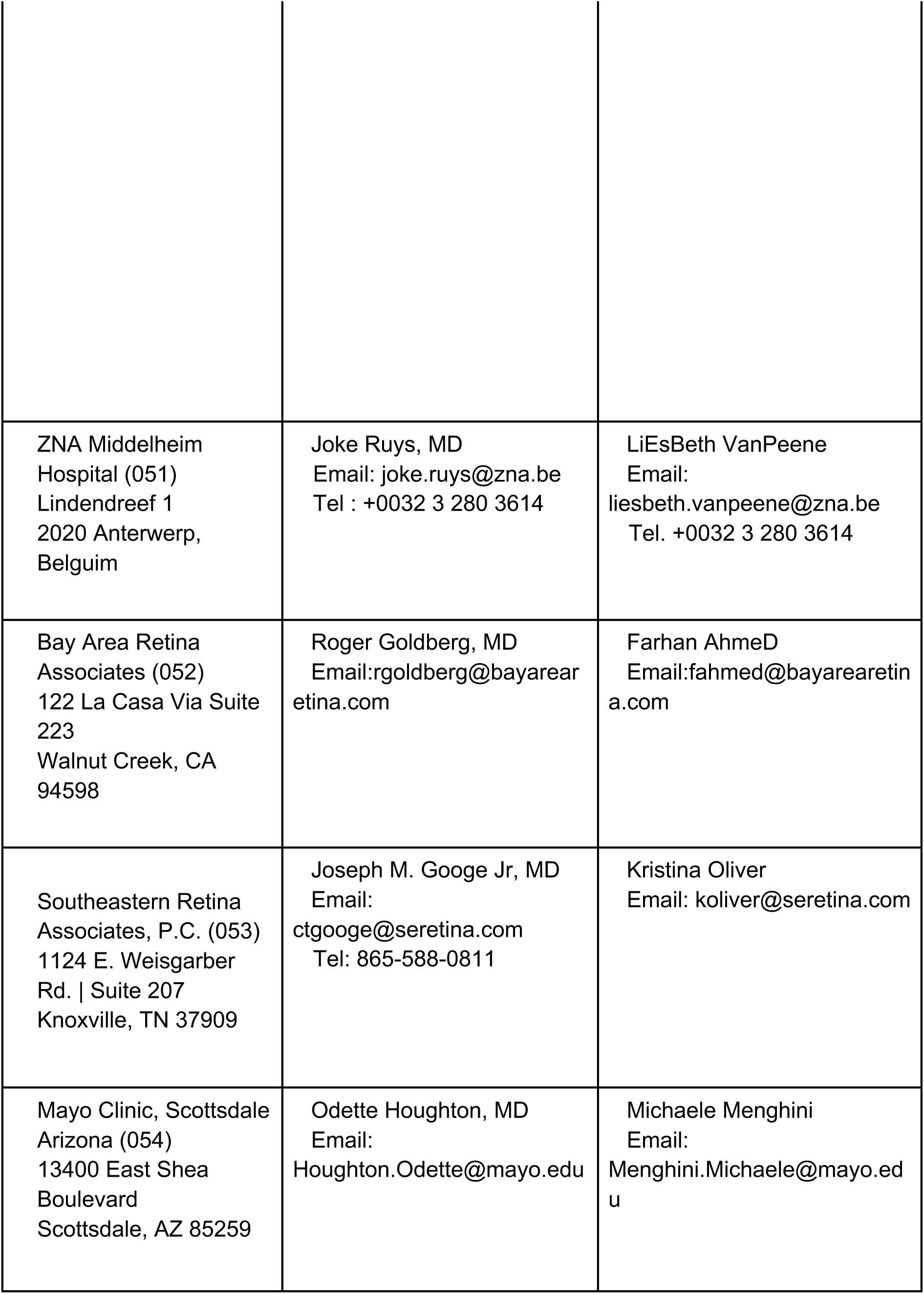
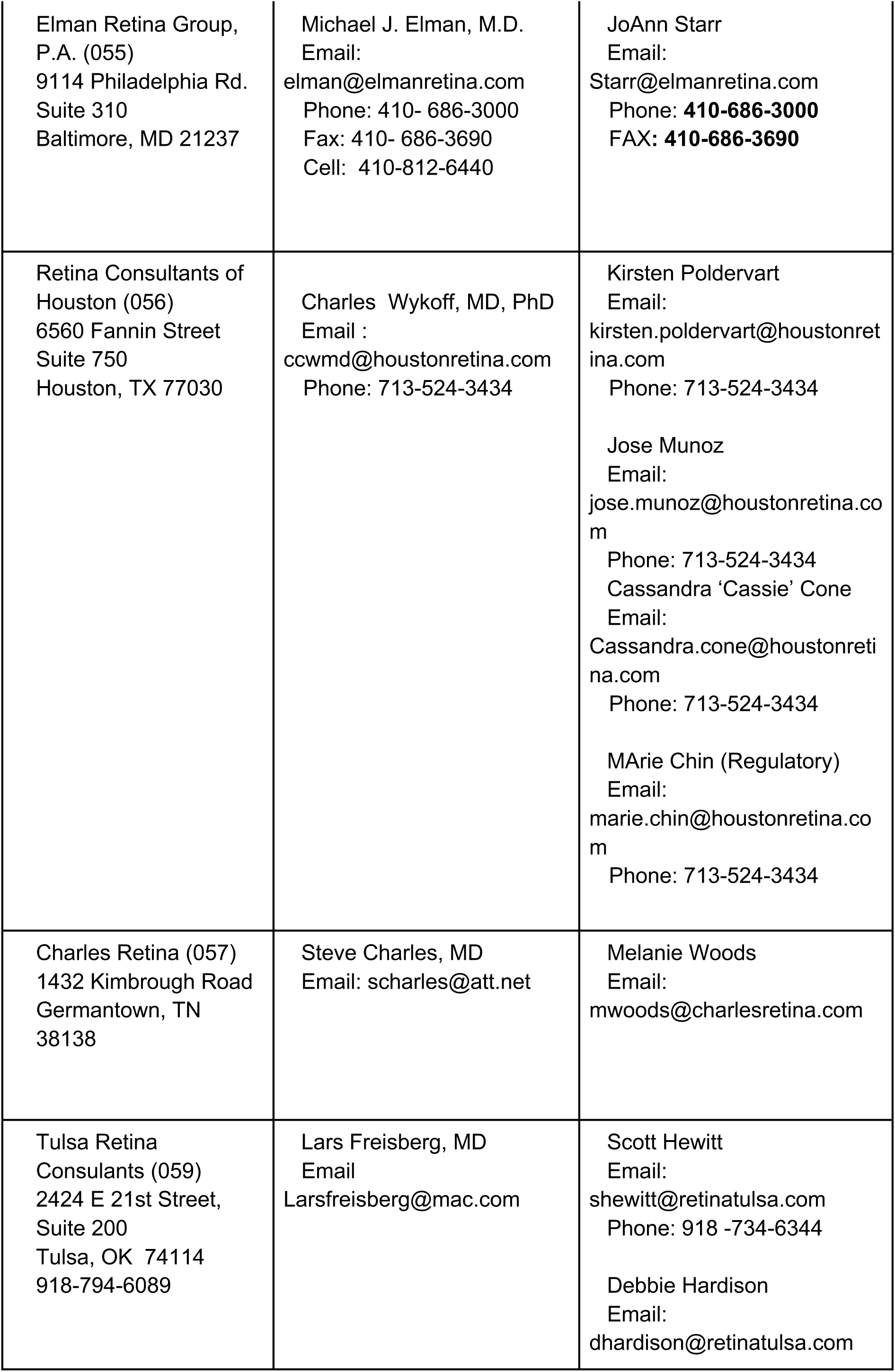
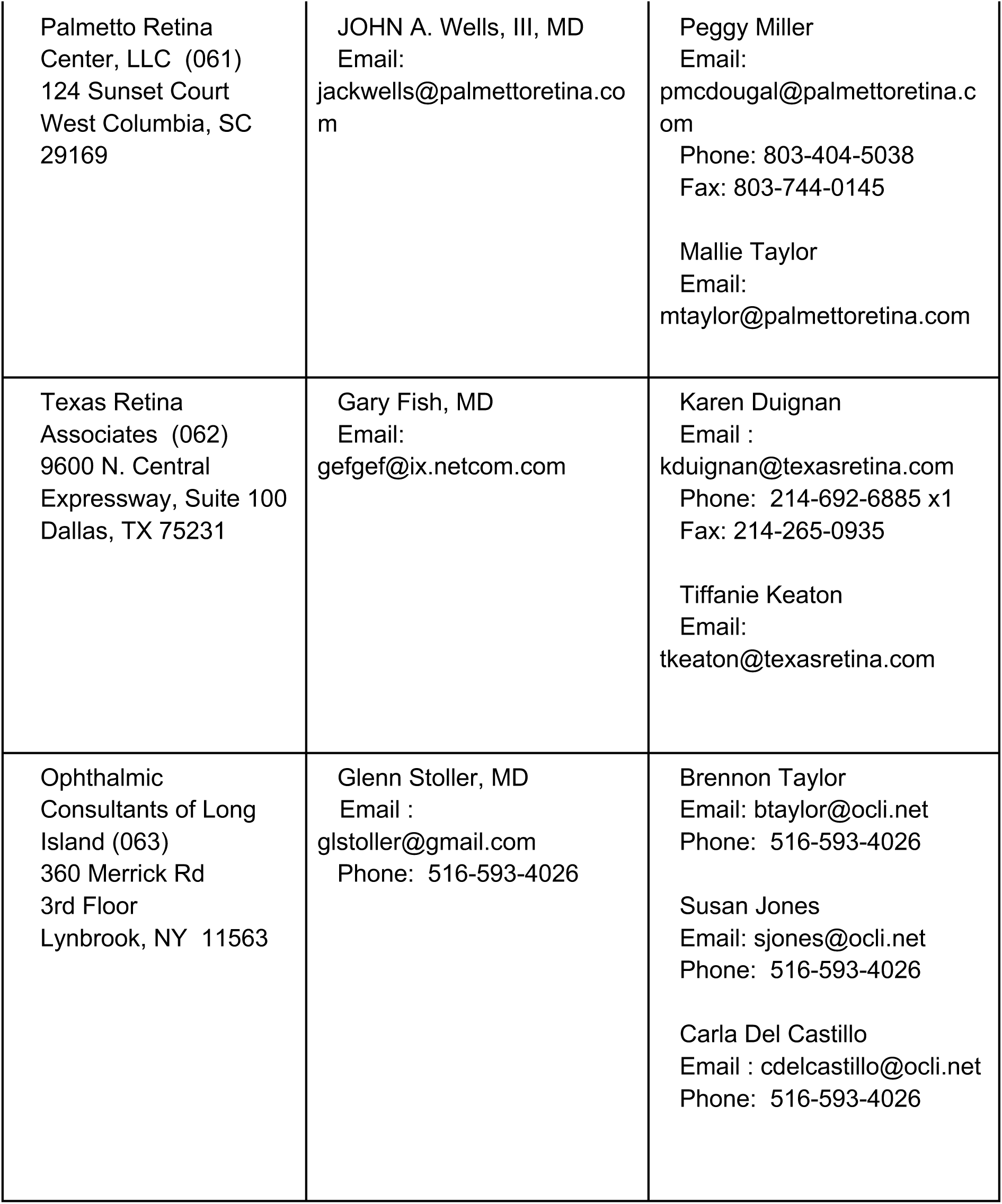
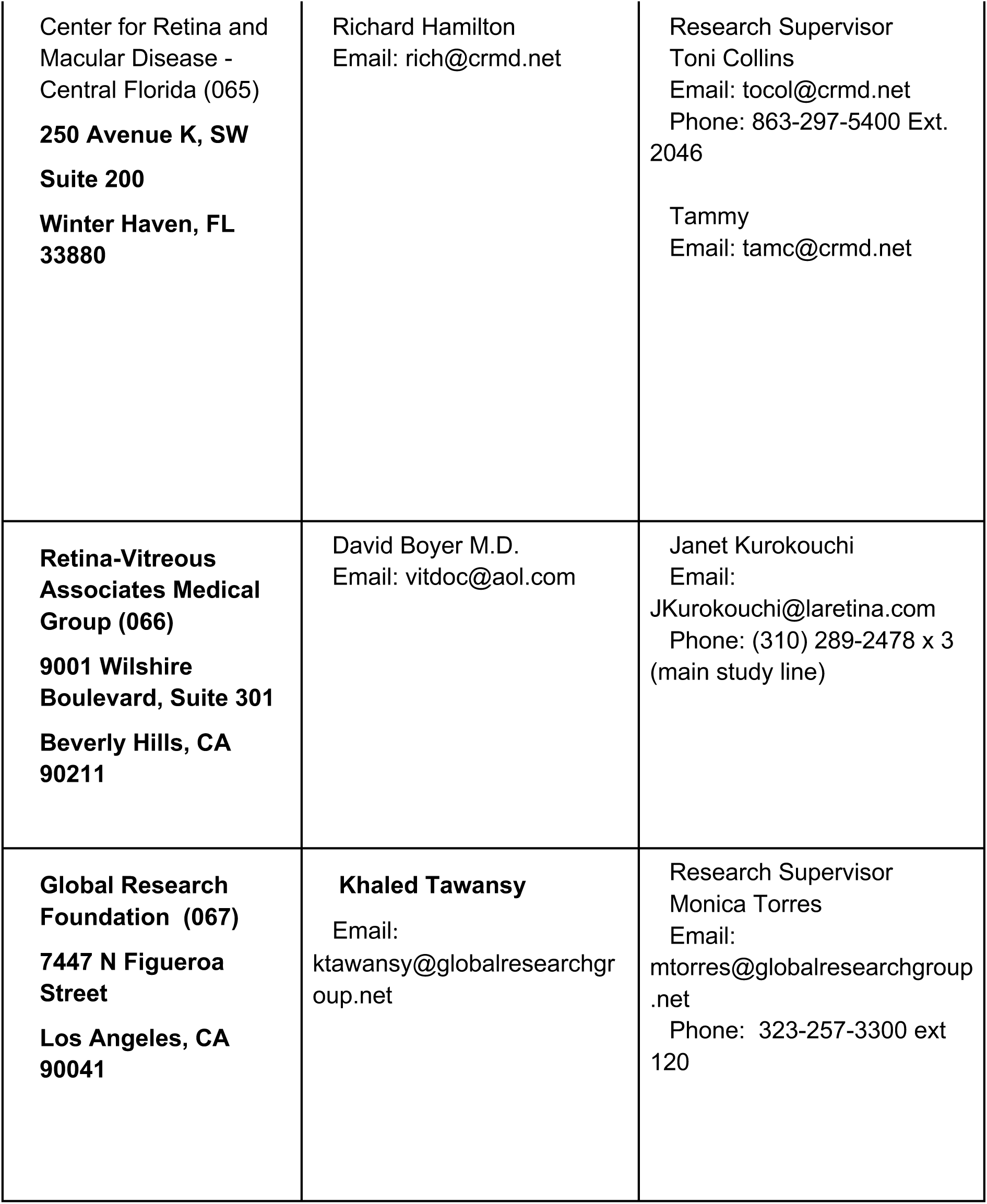
Supplementary Table 8:

